# Widespread haploid-biased gene expression in mammalian spermatogenesis associated with frequent selective sweeps and evolutionary conflict

**DOI:** 10.1101/846253

**Authors:** Kunal Bhutani, Katherine Stansifer, Simina Ticau, Lazar Bojic, Chloe Villani, Joanna Slisz, Claudia Cremers, Christian Roy, Jerry Donovan, Brian Fiske, Robin Friedman

## Abstract

1

Mendel’s first law dictates that alleles segregate randomly during meiosis and are distributed to offspring with equal frequency, requiring sperm to be functionally independent of their genetic payload. Developing mammalian spermatids have been thought to accomplish this by freely sharing RNA from virtually all genes through cytoplasmic bridges, equalizing allelic gene expression across different genotypes. Applying single cell RNA sequencing to developing spermatids, we identify a large class of mammalian genes whose allelic expression ratio is informative of the haploid genotype, which we call genoinformative markers (GIMs). 29% of spermatid-expressed genes in mice and 47% in non-human primates are not uniformly shared, and instead show a confident allelic expression bias of at least 2-fold towards the haploid genotype. This property of GIMs was significantly conserved between individuals and between rodents and primates. Consistent with the interpretation of specific RNA localization resulting in incomplete sharing through cytoplasmic bridges, we observe a strong depletion of GIM transcripts from chromatoid bodies, structures involved in shuttling RNA across cytoplasmic bridges, and an enrichment for 3’ UTR motifs involved in RNA localization. If GIMs are translated and functional in the context of fertility, they would be able to violate Mendel’s first law, leading to selective sweeps through a population. Indeed, we show that GIMs are enriched for signatures of positive selection, accounting for dozens of recent mouse, human, and primate selective sweeps. Intense selection at the sperm level risks evolutionary conflict between germline and somatic function, and GIMs show evidence of avoiding this conflict by exhibiting more testis-specific gene expression, paralogs, and isoforms than expression-matched control genes. The widespread existence of GIMs suggests that selective forces acting at the level of individual mammalian sperm are much more frequent than commonly believed.

**Author’s summary:** Mendel’s first law dictates that alleles are distributed to offspring with equal frequency, requiring sperm carrying different genetics to be functionally equivalent. Despite a small number of known exceptions to this, it is widely believed that sharing of gene products through cytoplasmic bridges erases virtually all differences between haploid sperm. Here, we show that a large class of mammalian genes are not completely shared across these bridges, therefore causing sperm phenotype to correspond partly to haploid genotype. We term these genes “genoinformative markers” (GIMs) and show that their identity tends to be conserved from rodents to primates. Because some GIMs can link sperm genotype to function, they can be thought of as selfish genetic elements which lead to natural selection between sperm rather than between organisms, a violation of Mendel’s first law. We find evidence of this biased inheritance, showing that GIMs are strongly enriched for selective sweeps that spread alleles through mouse and human populations. For genes expressed both in sperm and in somatic tissues, this can cause a conflict because optimizing gene function for sperm may be detrimental to its other functions. We show that there is evolutionary pressure to avoid this conflict, as GIMs are strongly enriched for testis-specific gene expression, testis-specific paralogs, and testis-specific isoforms. Therefore, GIMs and sperm-level natural selection may provide an elegant explanation for the peculiarity of testis gene expression patterns, which are an extreme outlier relative to all other tissues.

## 3 Introduction

In diploid organisms, Mendel’s First Law dictates equal transmission of alleles to the next generation, with strong selective pressure maintaining this 50:50 ratio (Crow 1979). In mammalian spermatogenesis, a long stage of haploid development raises the possibility of allele-biased gene expression and extensive functional variation between mature sperm (Immler 2008). This could be deleterious, for example for important gene products encoded on the X chromosome that would be missing from Y-bearing sperm. However, haploid sperm precursors are equipped with a mechanism for sharing of gene products: cytoplasmic bridges connecting neighboring cells (Braun et al. 1989). Therefore, mature mammalian sperm are thought to be functionally diploid with very rare exceptions.

Most examples of transmission ratio distortion (TRD), i.e. known exceptions to Mendelian inheritance, are attributable to factors other than sperm heterogeneity. However, a handful of sperm functional differences linked to genotype have been reported. The mouse *t* haplotype, a selfish genetic element transmitted at a rate of up to 99% from heterozygotes, is the best understood case. The mechanism for its TRD is post-meiotic expression and a lack of sharing of *t complex responder* gene products across cytoplasmic bridges, resulting in differential motility (Véron et al. 2009). Likewise, *Spam1* gene products have been shown to be retained in haploid spermatids, underlying TRD in mice carrying certain Robertsonian translocations (Zheng, Deng, and P. Martin-DeLeon 2001). In a mouse model for Niemann-Pick disease, heterozygous knockouts of *Smpd1* have sperm with functional differences in mitochondrial membrane potential associated with their genotype (Butler et al. 2007). Recently, TLR7/8 inhibitors have been reported to differentially affect sperm with the X or Y chromosome (Umehara, Tsujita, and Shimada 2019). Nevertheless, it is widely assumed that most gene products are shared between mammalian gametes, erasing any allelic expression bias.

If, however, sperm functional variation were linked to genotype more often than commonly believed, it might provide an elegant explanation for some peculiar evolutionary phenomena. Testes and spermatids in particular are extreme evolutionary outliers, having far more unique tissue-specific expression patterns, tissue-specific paralogs, alternative isoforms, and selective sweeps compared to other tissues (Kleene 2005). Sexual selection and intragenomic conflict is often invoked to explain this bias, but haploid selection on genes with transmission ratio distortion could easily have contributed (Joseph and Kirkpatrick 2004). For example, alleles with beneficial effects in mature sperm might have deleterious effects in somatic cells, which could drive avoidance of this conflict by evolving sperm-specific paralogs or isoforms. Widespread transmission ratio distortion would be difficult to observe directly due to rapid fixation of beneficial alleles and depletion of deleterious ones, but might leave traces over evolutionary timescales, altering the properties of testis-expressed genes.

TRD enabled by retention of haploid gene products in spermatids would require specific RNA localization rather than free diffusion across cytoplasmic bridges. Recent methodological advances in RNA detection have revealed widespread asymmetric mRNA distributions in a wide variety of cell types, including up to 70% of mRNAs during *D. melangogaster* development (Lécuyer et al. 2007; Buxbaum, Haimovich, and Singer 2015).

We therefore hypothesized that many endogenous mRNAs would be transcribed in haploid spermatids and incompletely shared across cytoplasmic bridges, resulting in allelic expression bias correlating to the sperm genotype (Fig. 1A). Since mature sperm are transcriptionally and translationally silent, allelic biases in mature sperm protein correlated with the haploid genotype would have to correspond to mRNA expression biases at the haploid spermatid stage. We therefore performed single cell RNA sequencing in spermatids (Fig. 1B) from hybrid mice and cynomolgus macaques, quantifying allele-specific biases in expression. We found surprisingly widespread chromosome-scale biases in single cells allowing confident identification of genes with strong allelic expression links to the genotype, which we term <u>g</u>eno<u>i</u>nformative <u>m</u>arkers (GIMs). We show evidence for subcellular localization patterns that help explain their lack of sharing across cytoplasmic bridges, as well as evolutionary consequences consistent with sperm-level natural selection.

**Figure 1:**
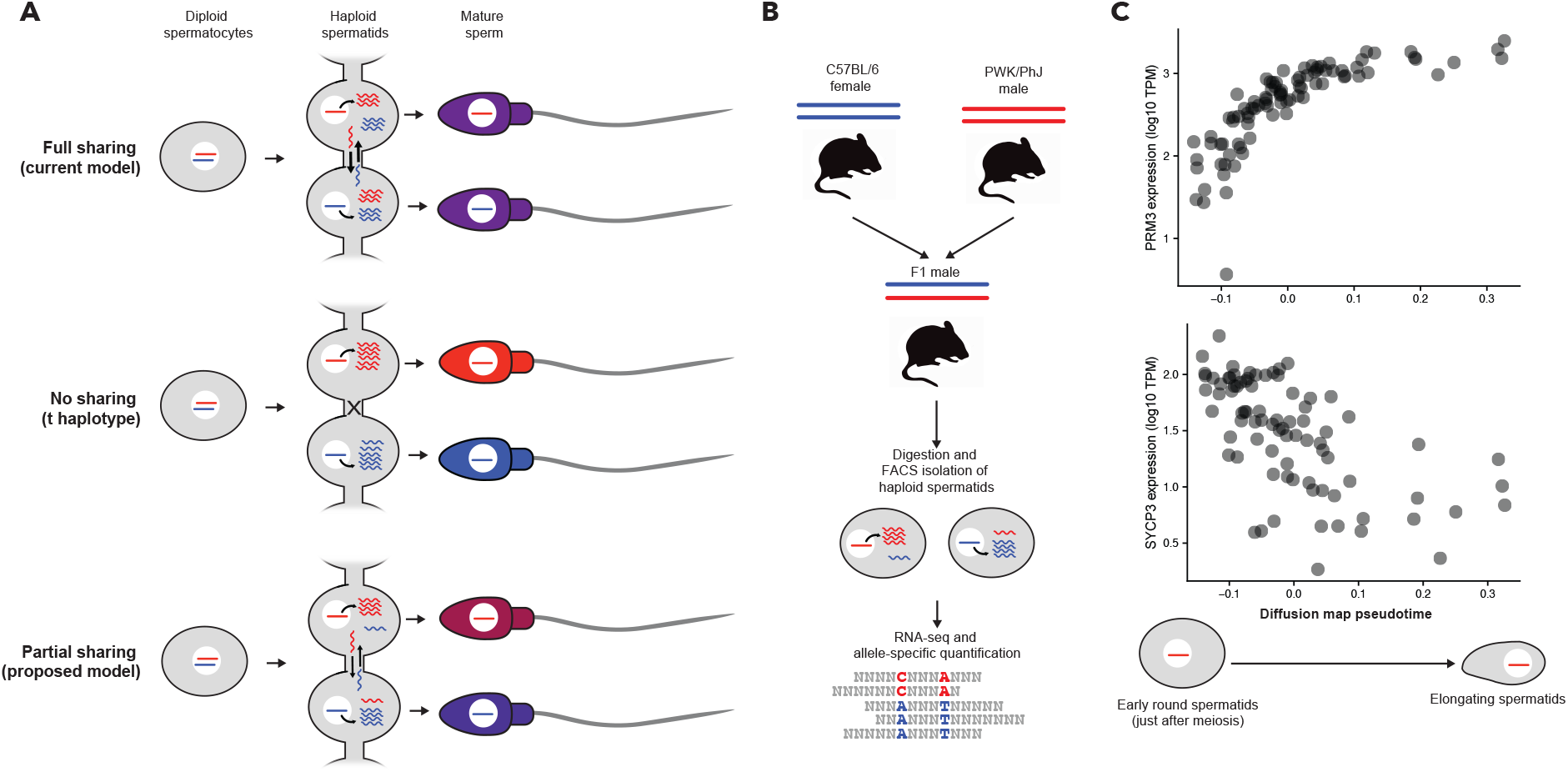
Single cell sequencing of haploid spermatids for assessing allelic bias. **A)** Models for allelic expression bias informative of the haploid genotype (genoinformative expression). The null hypothesis predicts complete sharing between spermatids, erasing any systematic allelic expression differences in mature sperm (top). Selfish genetic elements like the mouse *t haplotype* have virtually no sharing and lead to dramatic allelic differences in mature sperm (center), but incomplete sharing of transcripts would also lead to genoinformative expression (bottom). DNA is represented as straight lines with color representing an allele, and RNA is represented as wavy lines. Sperm color represents the degree of functional links to the allelic genotype. **B)** Experimental setup for single cell RNAseq. We crossed distantly related inbred mouse strains, digested single cells from the testis and enriched for haploid spermatids, and performed full-length RNA-seq and allele-specific quantification. **C)** Pseudotime analysis shows haploid spermatids covered a range from the early round stage (low expression of protamines) to the late elongating phase (very low expression of *SYCP3*)

## 4 Results

### 4.1 Many genes have allelic expression bias reflecting the haploid genotype in spermatids

We first set out to identify cases of incomplete sharing of RNA across cytoplasmic bridges in haploid spermatids (Fig. 1A). This would result in shared information (i.e. correlation) between the allelic expression of a gene and the haploid genotype of the cell, which we call genoinformative expression. Most single cell RNAseq experiments are poorly suited to quantifying allele-specific expression because they do not sequence samples from fully phased individuals, they only sequence a short tag from each RNA molecule (which may not contain a heterozygous site), and they do so with relatively low capture efficiency. To maximize the accuracy of our allele-specific quantification, we used an F1 hybrid (therefore fully phased) of distantly-related inbred mouse models, C57BL/6 and PWK/PhJ, having over 20 million heterozygous SNPs, compared to roughly 3 million in a human genome (Fig. 1B). We digested testis tissue to isolate single cells from their cytoplasmic bridges, enriched for haploid cells by flow cytometry, and performed full-length single cell RNA sequencing using a slightly modified SmartSeq2 protocol optimized for sensitive RNA capture (Methods).

Of 144 cells obtained from a single male mouse having successful RNA amplification, 126 passed filters as likely singlets with substantial read counts. Principal Components Analysis (PCA) and t-Distributed Stochastic Neighbor Embedding (t-SNE) revealed a mixture of three cell types expressing marker genes for spermatids, spermatocytes, and spermatogonia, respectively (Fig. S1A-C). Focusing on the 95 haploid spermatids, we used diffusion mapping (Angerer et al. 2016) to define a pseudotime space covering their differentiation process. The pseudotime ranges from early round spermatids up until the point that the number of genes expressed decreases rapidly at the elongation stage, when transcription arrests (Fig 1C, Fig. S1D). Late spermatid markers such as *PRM3* increase in expression over this pseudotime, while spermatocyte markers such as *SYCP3* decrease (Fig. 1C).

10,991 genes passed filters for calculation of genoinformative expression, including having at least one heterozygous site and having comparable mean expression of each allele (see Methods). We first focused on autosomes rather than sex chromosomes, because we could use the two alleles as an internal control, yielding an easily quantifiable allelic expression ratio within each cell. Visualizing allelic expression in individual haploid cells, we observed strong biases across large stretches of chromosomes, but no consistent bias in diploid controls (Fig. 2A, S2A). Across all haploid autosomes, there was a significant correlation of allelic ratios between neighboring genes that gradually decreased with chromosomal distance, and this correlation was completely absent in diploid controls (Fig. S1E-F). We reasoned that this effect could be explained by a combination of correlation caused by widespread genoinformative expression and degradation of this correlation with distance by recombination. Therefore, we designed a Bayesian probability framework based on an extension of a Hidden Markov Model to infer the haploid genotype of each cell including recombination breakpoints jointly with genoinformativity. Genoinformative expression was modeled as emissions based on the underlying genotype and propensity of an RNA to be shared across cytoplasmic bridges. Intuitively, this model shares information between genes across an entire chromosome for each cell, which means that even weak and noisy genoinformative expression signals in individual genes can aggregate to yield robust signals across large stretches of a chromosome. The model output a probability of genotypes for each cell, and a genoinformativity score for each gene representing the estimated fraction of transcripts retained from its haploid gene expression. Visual inspection confirmed that our inferred genotypes matched the observed expression biases well (Fig. 2A, Fig. S2A). If the inferred genotypes are accurate, the distribution of recombination breakpoints should follow the known recombination density in the mouse genome. Indeed, we saw a significant correlation of inferred recombination density to the published map (Cox et al. 2009) with good agreement at a resolution of 10 to 20 megabases (Fig 2B, S2B-C).

**Figure 2:**
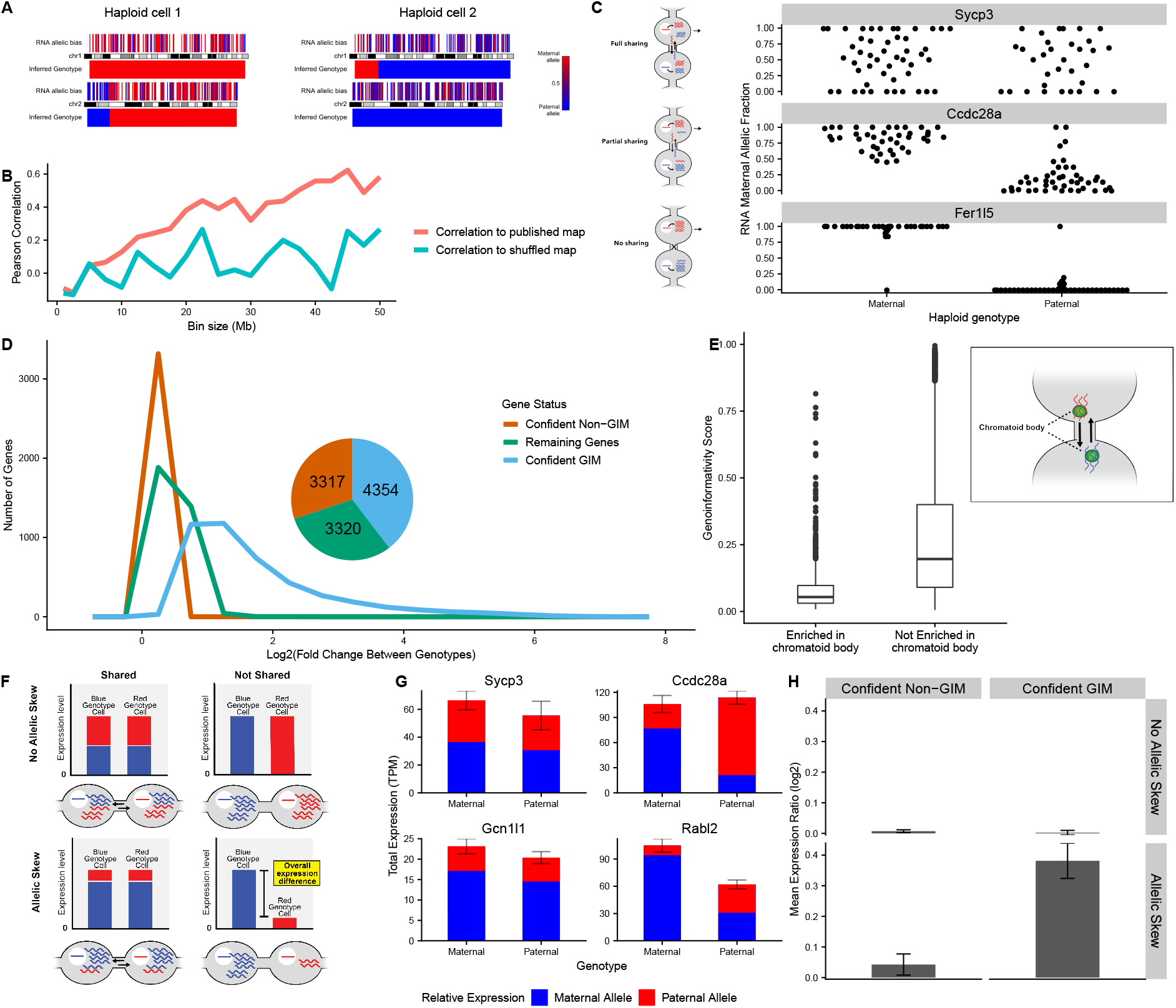
A large fraction of mouse genes exhibit genoinformative expression. **A)** Visualization of allelic bias in the first two chromosomes of two representative haploid cells. Each expressed gene is represented as a vertical line with color representing its allelic ratio (red for more maternal allele, blue for more paternal). Below each chromosome is the genotype inferred by our Bayesian method. **B)** Correlations between inferred recombination densities and a published mouse recombination map (Cox et al. 2009) or a control with recombination densities shuffled between all bins. As bin sizes decrease below about 20 megabases, the variance in our inferred rates increases, causing a degradation of our signal to noise ratio. **C)** Example genes illustrating differing levels of genoinformative expression (right) with their models of sharing (left). *Sycp3* exhibits no association with the haploid genotype, *Ccdc28a* exhibits a strong but incomplete association between the inferred genotype and the expressed allele, and *Fer1l5* exhibits a near-perfect correlation with the inferred genotype. **D)** GIM classification of all genes. Histogram shows the log2 of the expression ratio between the concordant allele (i.e. matching the genotype) over the discordant allele on average across cells. Inset: the total number of genes classified in each category of genoinformative expression. **E)** Genes with mRNAs enriched in the chromatoid body have significantly lower genoinformativity scores. Genoinformativity scores range from zero to one and represent the estimated fraction of transcripts originating from a cell’s haploid transcription. Inset: depiction of the chromatoid body’s role in shuttling mRNAs across cytoplasmic bridges in haploid spermatids. **F)** A model for how allelic skew (e.g. due to eQTLs) interacts with genoinformative expression. Only genes with both allelic skew and genoinformative expression (not shared) have their mean expression level correlated to the haploid genotype. **G)** Example genes matching the categories in (F). Only *Rabl2* has a significant mean expression difference (*p* = 1.5 × 10^−5^, Wilcoxon test). **H)** Summary of expression differences (log2 ratio of genotype concordant with skew to discordant) in all genes in each of the four combinations listed. Only with both allelic skew and GIMs is there an expression difference between cells of differing genotypes.

Examining for individual genes the concordance between allelic expression and haploid genotype across cells, we observed a wide range of genoinformativity (Fig. 2C): Many genes, like *Sycp3*, had no association between their allelic expression ratio and the inferred genotype, consistent with our null hypothesis of complete sharing across cytoplasmic bridges erasing allelic expression differences; some, such as *Fer1l5*, had virtually complete concordance with their inferred genotype, suggesting minimal sharing across cytoplasmic bridges; a larger set of genes had clear but intermediate genoinformativity, exemplified by *Ccdc28a*, suggesting partial sharing through cytoplasmic bridges. To determine thresholds for confident genoinformativity, we ran our Bayesian algorithm on shuffled data to create an empirical background expectation under the null hypothesis of no genoinformative expression (Fig. S2D-E). Thresholds of parameters for both the posterior distribution of the genoinformativity score and the strength of haplotype inference were selected to achieve an empirical False Discovery Rate of 10%. For convenience, genes that met the criteria for confident genoinformative expression were called <u>g</u>eno<u>i</u>nformative <u>m</u>arkers (GIMs), regardless of their effect size. Of the 10,991 genes for which we could estimate genoinformativity, 4,354 (39.6%) were confident GIMs and 3,317 (30.2%) were confidently not GIMs (see Methods; Fig. 2D, inset). We were unable to make a confident call for the remaining 3,320 (30.2%) due to marginal signal for genoinformativity. Of the confident genoinformative set, a wide range of effect sizes was seen, but 3,159 (28.8%) had at least a 2-fold average allelic expression ratio in favor of the allele matching the haploid genotype (Fig. 2D).

We were surprised that as many as a third of genes were classified as strong GIMs, so we sought to confirm our assumption that this corresponded to incomplete sharing across cytoplasmic bridges. The chromatoid body is a membraneless organelle (a phase-separated condensate) in germ cells that has been shown to shuttle RNA across cytoplasmic bridges to facilitate sharing (Fig. 2E inset; Ventelä, Toppari, and Parvinen 2003). We found that a published set of genes enriched in the chromatoid body (Meikar et al. 2014) had far lower genoinformativity scores than other genes (Fig. 2E), and that there were fewer GIMs enriched in the chromatoid body than expression-matched controls (Fig. S5C). This confirms that GIMs have different subcellular localization of their RNAs from non-GIMs.

### 4.2 GIMs have specific subcellular localization resulting in incomplete sharing across cytoplasmic bridges

To identify what mechanisms might be responsible for the differential localization of GIMs, we compared GIMs to non-GIM controls that were matched for expression across spermiogenesis as closely as possible (Fig. S5A, Table S3-4, Methods). Most eukaryotic mRNA localization is dictated by RNA-binding proteins via sequence motifs in 3’ UTRs (Andreassi and Riccio 2009), so we performed an enrichment analysis for known motifs of RNA-binding proteins that are expressed in spermatids. We identified 26 motifs significantly enriched in GIMs relative to controls, and zero significantly depleted in GIMs (Table S5).

Similarly, a gene ontology enrichment analysis identified strong enrichment for GIMs for specific protein localizations, especially membrane associations and axoneme or other tail localizations (Table S6). To further refine this result, we performed an enrichment analysis with a comprehensive localization database (Binder et al. 2014). This revealed a strong enrichment for genes with annotated localization in neurons, including both dendrites and axons (Table S7), probably reflecting the fact that subcellular RNA localization has been best studied in neurons but is governed by principles applicable across cell types (Ryder and Lerit 2018). Together, these data suggest a mechanism for genoinformativity whereby RNA-binding proteins bring some mRNAs to specific subcellular locations distal from chromatoid bodies, thus partially avoiding sharing across cytoplasmic bridges.

As independent confirmation of our incomplete sharing model for GIMs, we sought to use the much larger set of RNAseq reads that did not overlap a heterozygous site but could be used for estimating overall expression levels. GIMs have allelic expression biases based on the haploid genotype, but because 50% of cells have each genotype, GIMs do not necessarily have a mean allelic expression bias when averaging across many cells (here called allelic skew). However, many genes have a mean allelic skew for other reasons, for example due to expression quantitative trait loci (eQTLs) wherein a genetic variant has differential effects on the expression of a gene. The incomplete sharing model predicts that genes may have different expression levels in spermatids with the paternal versus maternal genotype, but only when they have both an allelic skew and genoinformative expression (Fig. 2F). To illustrate this point, *Sycp3* (Non-GIM, no allelic skew), *Ccdc28a* (GIM, no allelic skew), and *Gcn1l1* (Non-GIM, 2.7-fold allelic skew) all have no difference in mean total expression from the maternal and paternal genotype cells (Fig. 2G). However, *Rabl2*, which has a 3.0-fold allelic skew and genoinformativity score of 0.45 has a significant difference in expression between the two spermatid genotypes (*p* = 1.2 × 10^−5^, t test). Across all genes, we observe that the expression level of GIMs with allelic skew is linked to the haploid genotype in the expected direction, but not for non-GIMs and not for genes without overall allelic skew (Fig. 2H). Therefore both allele-informative and non-allele-informative RNAseq reads support the identity of GIMs and the incomplete sharing model.

### 4.3 Sex chromosome genes also exhibit genoinformative expression

Although our Bayesian method for inferring genotype and genoinformativity cannot be applied to sex chromosomes due to the lack of allelic expression data, genoinformative expression of sex chromosome genes would provide an elegant explanation for models of sex ratio distortion in mice (Cocquet et al. 2012; Eep, Pji, and Ellis Email n.d.). We therefore developed a separate method to identify sex chromosome GIMs based on variation in expression levels rather than in allelic ratios. We started by reasoning that X chromosome GIMs should have correlated expression and be anticorrelated with Y GIMs. Because expression levels in any given spermatid can be strongly influenced by developmental stage, we first corrected for the position in the diffusion map pseudotime. Clustering genes by pairwise correlation after correction, we identified two distinct clusters that corresponded overwhelmingly to the X and Y chromosome, respectively (Fig. S3A). In contrast, performing the same analysis on autosomal controls yielded no similar clusters (Fig. S3B). We selected putative GIMs from these distinct clusters that displayed strong correlation signals (see methods), resulting in 63 X GIMs and 84 Y GIMs (Table S2). Spermatids tend to have high or low mean levels of X GIMs, but not intermediate levels (Fig. S3C). Therefore, sex chromosomes appear to be no exception to the prevalence of genoinformative expression, at least on a quantitative level.

### 4.4 Genoinformativity is conserved between individuals and across species

So far, we have only considered mice with one genetic background, so we next asked whether the phenomenon of widespread genoinformative expression extends to other mammals. We dissociated testes from two outbred cynomolgus primates (*Macaca fascicularis*), isolated haploid spermatids and performed single cell RNAseq. Cynomolgus monkeys have the advantage of being highly heterozygous, with ~13 million heterozygous SNPs per individual, compared to ~3 million for humans. Because our method for inferring genotypes relies on sharing information across entire chromosomes, we required fully phased chromosomes to quantify genoinformative expression. We therefore combined two phasing methods: a dense, short-range phasing using linked read sequencing, and a sparse, long-range phasing using whole genome sequencing of single haploid spermatids (Fig. 3A). Combining the two sources of information led to densely phased chromosomes for each individual, resulting in 11,654,918 and 10,131,178 phased sites in Cynomolgus 1 and 2, respectively (Fig. S4A).

**Figure 3:**
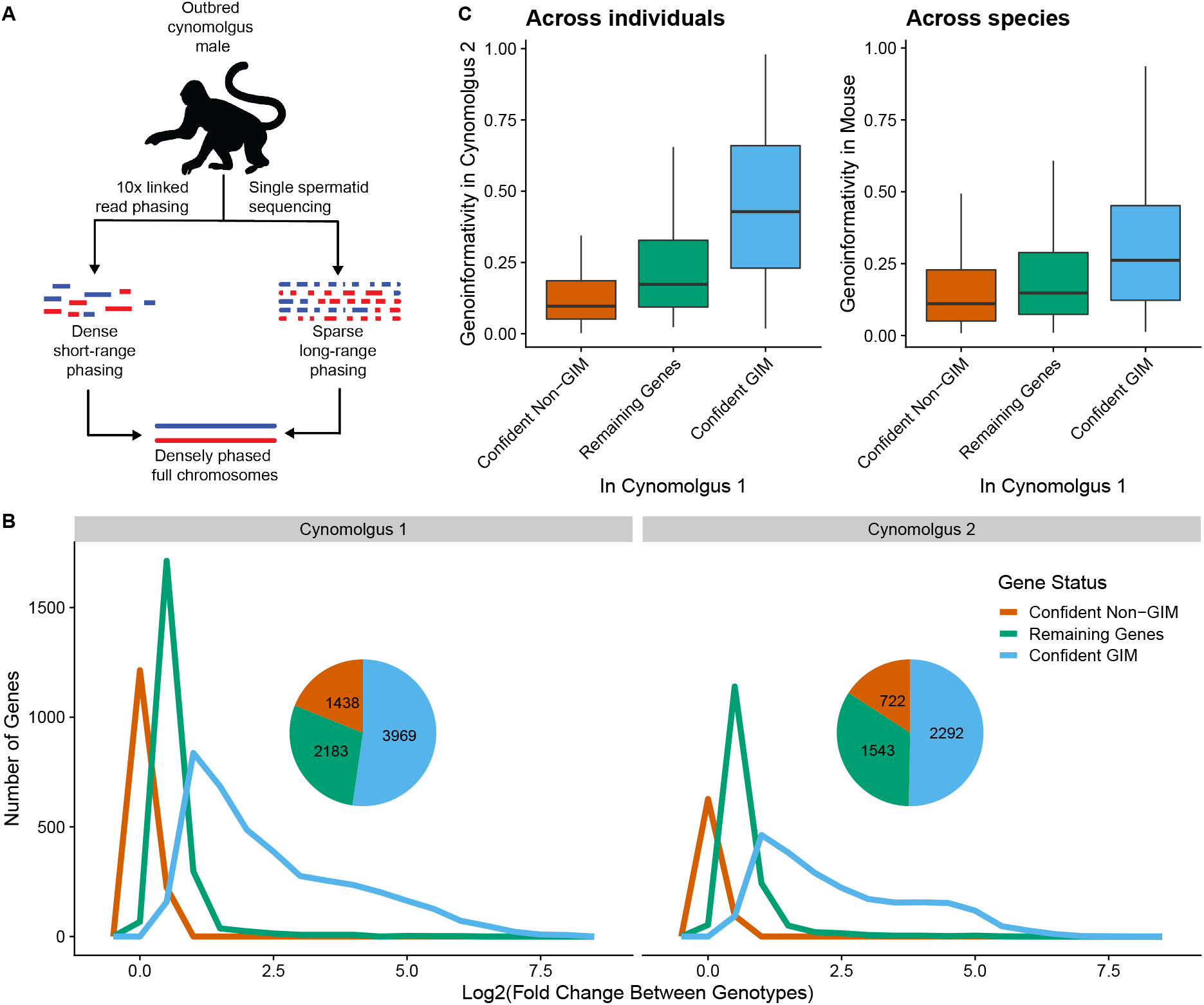
GIMs are conserved between individuals and across species. **A)** Fully phased chromosomes were generated directly from outbred cynomolgus individuals by computationally merging phasing maps from two experimental techniques: short-range phasing from 10x Genomics linked read sequencing, and long-range phasing from whole genome sequencing of several single haploid spermatids. **B)** Genoinformative expression classification of all genes as in Fig. 2D, for each of two cynomolgus individuals. Histogram shows the log2 of the expression ratio between the concordant allele and the discordant allele on average, where the concordant allele matches the inferred genotype. Inset: the total number of genes classified in each category of genoinformative expression. **C)** Conservation of genoinformativity. Genes are categorized based on their genoinformativity classification in Cynomolgus 1 (x axis), and genoinformativity is plotted for these genes in Cynomolgus 2 (left) or orthologs in mouse (right). Genoinformativity scores range from zero to one and reflect the degree of shared information with genotype.

We were able to quantify allelic expression of a smaller number of genes for cynomolgus spermatids than for mice (7,590 and 4,557 for the two cynomolgus compared to 10,991 in mice), mostly due a smaller number of heterozygous sites. Nevertheless, we observed comparable quality of our genotype inference, including significant correlation of inferred recombination rates between individuals, an expression skew in GIMs with allelic expression skew, and substantial differences between real and shuffled data (Fig. S4B-E). Again using an empirical false discovery rate of 10% in each individual, we classified 50.3% and 52.3% of spermatid-expressed genes as confident GIMs, respectively (Fig. 3B). The effect sizes were comparable to those seen in mice, with 44.6% and 43.3% of spermatid-expressed genes having at least a 2-fold average expression difference between alleles in favor of the haploid genotype. In total, 47.3% of genes that could be quantified met this threshold in either of the two individuals.

Because the two individuals have different heterozygous sites, only 2,366 genes had quantified genoinformativity in both. Among these genes, those that were classified as a confident GIM in one individual had far higher genoinformativity scores in the other individual, and those classified as a confident non-GIM had far lower genoinformativity scores in the other individual (*p* < 2.2 × 10^−16^; Fig. 3C). This suggests that within a species, the property of genoinformativity is highly consistent. To look across far larger evolutionary timescales, we compared cynomolgus genes to their orthologs in mouse with a genoinformativity score in each (n = 2,838). Confident GIMs in cynomolgus had higher genoinformativity in mouse than confident non-GIMs (*p* < 2.2 × 10^−16^; Fig. 3C), although the relationship was weaker than within a single species. This suggests that the features that confer incomplete sharing across cytoplasmic bridges evolve slowly, so that the identities of GIMs tend to be maintained across evolutionary timescales.

### 4.5 GIMs show signs of sperm-level natural selection and evolutionary conflict

The substantial fraction of genes having genoinformative expression at the RNA level is surprising, but it does not necessarily imply functional differences in sperm. For example, proteins could be shared across cytoplasmic bridges, nullifying any allelic differences at the RNA level. In contrast, if GIMs lead to functional differences in sperm linked to their genotype, sperm-level natural selection could result in increased evolutionary forces (both purifying and positive selection) acting on GIMs compared to other genes. Given that the identities of GIMs have been maintained across an appreciable evolutionary distance, we reasoned that functional differences in GIMs would lead to detectable signatures in the genome even if they rarely arise. Selective sweeps entail a beneficial allele experiencing positive selection and rapidly reaching fixation in a population, which leaves a signal that can be detected by a variety of statistical tests over patterns of variation in the genome. We cross-referenced a set of selective sweeps in wild mouse populations (Staubach et al. 2012) with GIMs and non-GIM controls, either randomly selected from spermatid-expressed genes or matched for expression patterns across spermiogenesis. The GIMs were found in significantly more selective sweep regions than expected by chance (*p* = 3 × 10^−25^) corresponding to an excess of 47 ± 4.6 selective sweeps putatively attributable to genoinformativity (Fig 4A, left). Although we do not know of studies of selective sweeps in cynomolgus, we took advantage of abundant predictions of selective sweeps in humans by examining orthologs of cynomolgus GIMs and non-GIMs. Using a set of human selective sweeps (Refoyo-Martínez et al. 2019), we find a significant enrichment of GIMs (*p* ≤ .013) corresponding to 9.4 ± 4.2 sweeps putatively attributable to genoinformativity (Fig 4A, right). We corroborated this enrichment for GIMs in a wide variety of tests for selective sweeps in humans and primates on multiple timescales (Fig. S5B). Examining an even larger set of tests for natural selection using 1000 genomes project data (Pybus et al. 2014), we found significant enrichments in a majority of tests (Fig. S5D). Together, this indicates that GIMs are associated with an increased rate of positive selection over evolutionary time.

**Figure 4:**
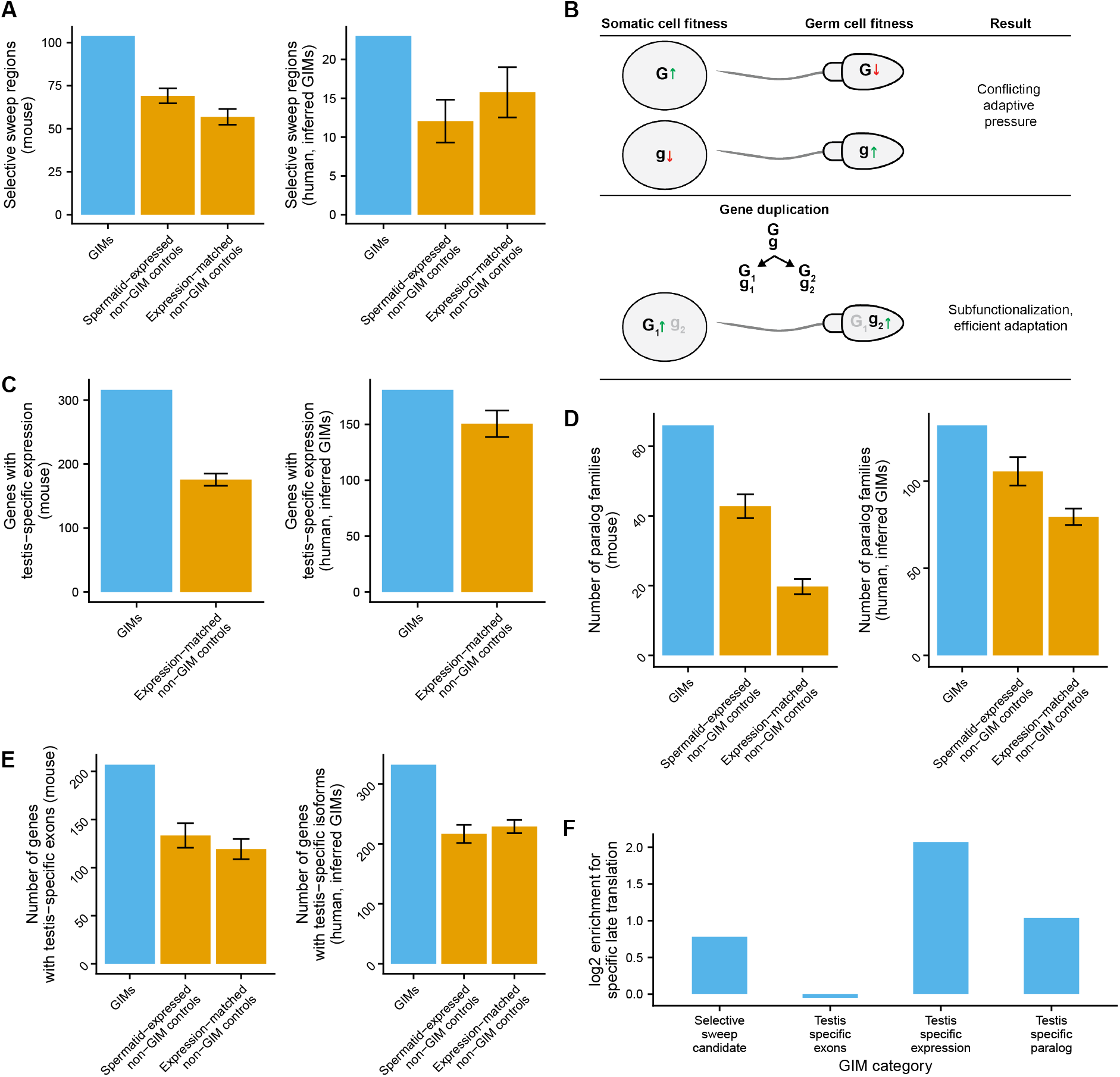
GIMs are associated with sperm-level natural selection and evolutionary conflict. **A)** GIMs are enriched in selective sweep regions in mouse (Staubach et al. 2012) and human (Refoyo-Martínez et al. 2019). Human GIMs were inferred from cynomolgus orthologs. GIMs were compared to control sets (orange bars), either selected from all spermatid-expressed confident non-GIMs, or confident non-GIMs matched to GIMs by their spermatid expression trajectory. **B)** Model for evolutionary conflict between sperm-level and organism-level natural selection. The gene has one allele with beneficial effect in somatic cells but detrimental effect in sperm (*G*) and one allele with the reverse pattern (*g*), resulting in positive selection for *g* at the sperm level, but negative selection at the organism level. A resolution to conflict can be achieved by duplication into two genes, *G*_1_/*g*_1_ expressed in somatic cells and *G*_2_/*g*_2_ expressed in sperm. Selection will then favor the *G*_1_ and *g*_2_ alleles, with no detrimental effects at either level. **C)** GIMs are enriched for testis-specific expression in mice and human, defined as 10-fold higher expression than any other tissue. GIMs were only compared to non-GIMs matched for spermatid expression trajectory, because testis-specific expression is by definition dependent on spermatid expression level. **D)** GIMs represent a higher number of paralog families than non-GIMs in mice and humans. Controls as in (A). **E)** GIMs are enriched in testis-specific isoforms in humans and testis-specific exons in mice. Controls as in (A). **F)** GIMs that are functional candidates are enriched for specific late translation. The GIMs are taken from the blue bars in panel A, C, D, and E, respectively. GIMs in each functional category are compared with GIMs not in that category, and the proportion with specific late translation was calculated. The log2 of the ratio of these proportions is plotted.

**Figure 5:**
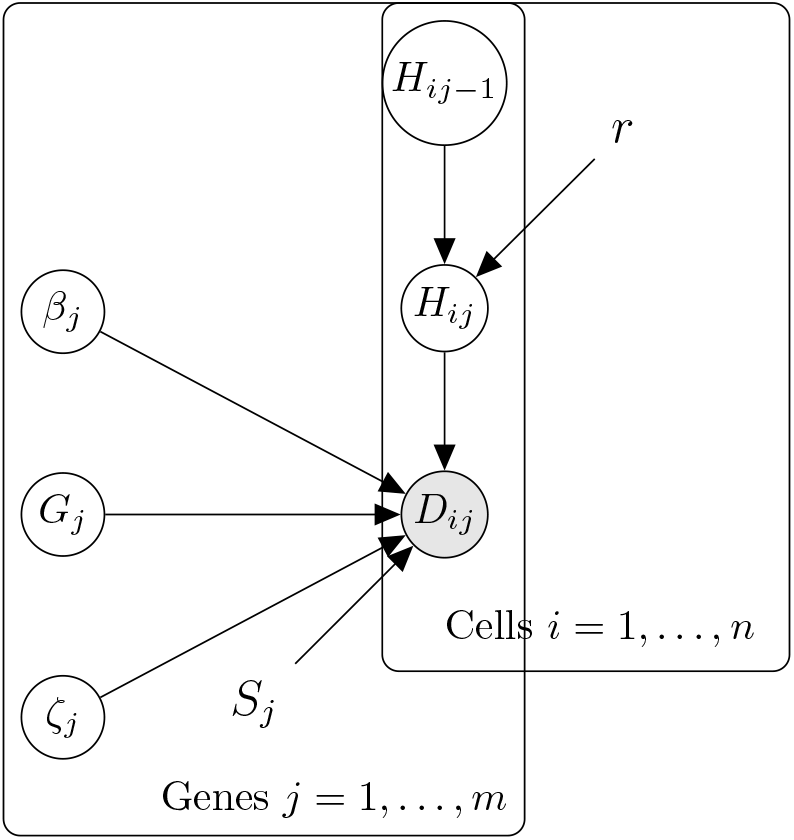
Graphical model for Bayesian method

Sperm-level natural selection poses an evolutionary conundrum: due to its highly specialized function, what is good for the sperm is not necessarily good for the organism. In other words, selection for a beneficial allele in sperm may decrease overall fitness if the allele is deleterious in a somatic cell context (Fig. 4B). Over evolutionary time, this conflict might make genoinformative expression deleterious for genes with somatic functions, but not for genes uniquely expressed in male reproductive tissue. Supporting this hypothesis, we see that GIMs are more likely to be testis-specific in both mouse (*p* < 10^−22^) and human (*p* = 0.006; Fig. 4C). When it arises, the evolutionary conflict caused by sperm-level selection will cause evolutionary pressure for separating functions for the gene in germ and somatic cells. Examples of this evolutionary pattern include gene duplication followed by subfunctionalization (Fig. 4B), and testis-specific gene isoforms. As predicted, GIMs are significantly enriched in paralog families that are predominantly testis-expressed in both mouse (*p* ≤ 6.7×10^−12^; Fig. 4D, left) and human (*p* ≤ 0.0007; Fig. 4D, right). Human GIMs are also enriched testis-specific isoforms (*p* ≤ 1.9 × 10^−14^; Fig. 4E, right), and although we are not aware of similar quality isoform-level mouse datasets, mouse GIMs are significantly more likely to have testis-specific exons (*p* ≤ 3.7 × 10^−9^; Fig. 4E, left).

Each of these lines of evidence implies that GIMs with these properties are enriched for causing functional differences in sperm, which would require incomplete sharing of proteins across cytoplasmic bridges. In the mouse *t haplotype*, this occurs in part by translating a protein late in spermiogenesis, as cytoplasmic bridges start to break down (Véron et al. 2009). We therefore predicted that GIMs enriched for causing functional differences in sperm would also be enriched in late translation of their proteins compared to other GIMs. Examining a polysome profiling dataset across mouse spermatogenesis (Iguchi, Tobias, and Hecht 2006), mouse GIMs that were functional candidates based on selective sweeps, testis-specific expression, or testis-specific paralogs, were indeed enriched for late translation (p = 0.045, 1.4 × 10^−12^, 0.00045, Fisher’s exact test; Fig. 4F). However, we did not see enrichment in late translation for GIMs that had testis-specific exons. These results suggest that late translation of GIMs is one mechanism by which they may lead to sperm-level functional differences, causing a higher rate of selective sweeps and avoidance of evolutionary conflict.

## 5 Discussion

Here we have shown that a large fraction of spermatid-expressed genes are not completely shared between haploid spermatids, resulting in allelic expression that is linked to the haploid genotype, which we call genoinformative expression. Our model for the mechanism for this genoinformative expression is subcellular localization of RNAs, occurring through RNA-binding protein motifs in the 3’ UTRs or other mechanisms, resulting in depletion of GIMs from the chromatoid body (which facilitates sharing across cytopasmic bridges). GIMs are substantially conserved across populations and evolutionary timescales, so we predict these mechansisms are conserved as well.

In light of this finding, a number of cases of sperm-level functional differences in the literature can be putatively attributed to GIMs (Conway et al. 1994; P. A. Martin-DeLeon et al. 2005; Butler et al. 2007; Véron et al. 2009; Cocquet et al. 2012; Alavioon et al. 2017; Nadeau 2017; Umehara, Tsujita, and Shimada 2019). Despite the growing number of examples of sperm-mediated transmission ratio distortion, it has been widely assumed these are isolated cases and that mammalian sperm are functionally diploid as a rule. The fact that GIMs were so common (over a third of spermatid-expressed genes) surprised us, and suggests that many more cases of sperm-level functional heterogeneity based on genotype will be found.

Mendel’s first law dictates that alleles of genes are inherited with equal probability, requiring sperm to be functionally equivalent regardless of their haploid genotype. We believe that remains the case for the majority of genes in mammals at any given time, since transmission ratio distortion has not been commonly observed. However, we show that over evolutionary timescales, GIMs are associated with an increased rate of selective sweeps, suggesting selection at the level of sperm based on functional differences linked to alleles. At first glance, reconciling the sperm-level selection with the predominance of Mendel’s first law seems difficult, but there are several reasons to believe they are compatible: 1) We find evidence for only tens to hundreds of selective sweeps across deep timescales and across thousands of GIMs, suggesting that they are relatively rare; 2) Selective sweeps happen quickly on an evolutionary timescale, erasing standing variation and making transmission ratio distortion a rare phenomenon at any one time; 3) Because most GIMs lead to only modest allelic differences (2-4 fold), sperm with these differences may be functionally equivalent or will lead to modest transmission ratio distortion, as is observed for example in mouse Yq deletions or Slx knockdowns (Conway et al. 1994; Cocquet et al. 2012), which is challenging to quantify in most mammals; and 4) Avoidance of evolutionary conflict by evolving sperm-specific expression removes cases of balancing selection, which might have resulted in observable transmission ratio distortion on standing variation.

While genoinformative expression is widespread at the RNA level, we do not have direct evidence for how common it is at the protein level. One reason to believe there are substantially fewer protein-level GIMs than RNA-level GIMs is that proteins can be shared across cytoplasmic bridges. This is consistent with the fact that GIMs that are preferentially translated late in spermiogenesis, when there is little to no time to be shared across cytoplasmic bridges, are enriched in evidence for selection or avoidance of evolutionary conflict. Even extremely late-expressed GIMs may not always lead to functional differences in sperm, because epididymal exosomes deliver proteins from diploid cells to sperm after they cease transcription and translation, potentially masking allelic differences in mature sperm. Another mechanism for masking the functional consequences of GIMs may be the abundant post-translational regulation of mature sperm, for example during capacitation, which might create larger cell-to-cell variation among sperm than GIMs.

Nevertheless, the ability of GIMs to lead to sperm-level natural selection may have profound evolutionary consequences. We have shown strong enrichments of GIMs for testis-specific expression, testis-specific paralogs, and testis-specific isoforms or exon usage. There are two forces that could give rise to these results: first, evolutionary conflict arises repeatedly in GIMs, which provides an evolutionary advantage to evolve distinct sperm-level function; second, that evolutionary conflict provides pressure to decrease genoinformativity (i.e. increase sharing across cytoplasmic bridges), so that the remaining GIMs preferentially have more sperm-specific expression. It is impossible to distinguish between these models with the data here, but it is likely that both forces contribute. More comprehensive catalogs of GIMs across species may be necessary to identify which is predominant.

This provokes a profound question: why, from an evolutionary perspective, do GIMs exist at all? For sex chromosome genes, such as *Akap4* (an X-chromosome gene required for sperm motility), it is clear that some degree of sharing is required for sperm function and specific mechanisms have evolved to facilitate sharing (Morales et al. 2002). However, it is not clear that genes need to be shared equally or that absolute functional equivalence is achieved; in some cases, a 2-fold or 4-fold difference in allelic expression may not have strong enough functional effects to exert evolutionary pressure to fully share transcripts. Also, sex chromosome genes are a special case that are hemizygous in males, so there may be even less pressure to share equally for autosomes. For genes or isoforms that are sperm specific, there could in fact be a benefit to sperm-level selection: an intensification of both purifying and positive selection by adding a selective layer on top of organism-level selection. In these cases, there would be no evolutionary conflict between the two selective layers, so some GIMs could become “selfish elements” whose interests are aligned with the organism: improving sperm function, which in turn increases the number of offspring.

The testis-expressed genome has long been a puzzling outlier, including by far the most tissue-specific gene expression, the most tissue-specific paralogs, and the most rapidly evolving genes. The widespread presence of GIMs raises the possibility that sperm-level selection and resulting evolutionary conflicts are common enough to provide an elegant explanation for these phenomena. If functional and molecular heterogeneity of sperm can be understood in enough detail, it is even possible that it could be exploited to isolate and eliminate sperm carrying severe Mendelian disease genes, reducing the risk of disease transmission across generations, as has been previously suggested (e.g. Butler et al. 2007). Given the rarity of GIM-related selective sweeps, it may be technically challenging to identify and leverage this expanded source of sperm heterogeneity. However, the surprisingly widespread existence of GIMs raises the possibility that a wide variety of severe diseases could be prevented by means of sperm selection.

## 6 Methods

### 6.1 Spermatid isolation and cell sorting

Testes were reduced to a single-cell suspension (breaking apart the intracellular bridges between germ cells in the process), using the two-step digestion protocol of Gaysinskaya et al. 2014. Digestions were performed in 6 ml for mouse, with one whole testis as starting material (tunica albuginea removed); and in 30 ml for non-human primate, with 600mg of diced testis tissue as starting material. First, to disperse the seminiferous tubules, testis tissue was incubated in digestion solution 1: Hanks’ Balanced Salt solution (HBSS, Sigma Aldrich), 1 mg/ml collagenase Type I (Worthington Biochemical), and 6 U/ml DNAse I (Sigma Aldrich). Incubation was at 37°C for 10 min with horizontal agitation. Tubules were then allowed to settle and the supernatant (containing somatic cells) was discarded. Digestion solution 2 was then added to reduce the tubules to a single-cell suspension: HBSS, 1 mg/ml collagenase, 6 U/ml DNAse, and 0.05% trypsin (Gibco, 2.5% stock solution). Incubation was for 25 min at 37°C with horizontal agitation; tubules were pipetted every 5 minutes, and an additional 0.025% trypsin was added halfway through the incubation. Successful digestion was confirmed by examining the cell suspension under a light microscope. Digestion was quenched with Newborn Calf Serum (Gibco).

After digestion, the single-cell suspension was filtered through a 100 μm cell strainer and centrifuged for 10 minutes at 500g. The supernatant was discarded, and the cell pellet was gently resuspended at 1-2×10^6^ cells/ml in PBS + 5 mg/ml BSA. Hoechst 33342 was added at 10 μg/ml and cells were incubated for 30 minutes at 37°C. Propidium iodide (PI) was added at 1 μg/ml during the last 5 minutes of incubation. Samples were filtered through a 40 μm mesh immediately before sorting.

Single live spermatids were then sorted into 96-well plates as described below, using a BD FACS Aria, a Beckman Coulter MoFlo Astrios, or a SONY Synergy SY3200 instrument. Our gating strategy was as follows: Selected for 1n cells (spermatids and sperm) based on Hoechst 33342 fluorescence intensity (with 355 nm excitation and a 448/59 nm bandpass emission filter) (Gaysinskaya et al. 2014); Selected for PI-negative cells to get a live population (PI was measured with 561 nm excitation and a 614/20 nm bandpass emission filter)); Enriched for round spermatids by selecting cells with high forward scatter (Bastos et al. 2005)

### 6.2 Cynomolgus primates

Adult male cynomolgus monkeys (*Macaca fascicularis*) were used for the non-human primate studies conducted at the University of Kentucky. Monkeys were singly housed in climate-controlled conditions with 12-hour light/dark cycles. Monkeys were provided water ad libitum and fed Teklad Global 20% Protein Primate Diet. Spermatid isolation and sorting was preformed at the University of Kentucky with two male monkeys. Monkeys were euthanized, testes were promptly removed and placed in Hanks’ Balanced Salt Solution (HBSS) on ice, prior to proceeding to tissue digestion and subsequent preparation of a single cell suspension for cell sorting. All animal care, procedures, and experiments were based on approved institutional protocols from the University of Kentucky Institutional Animal Care and Use Committee IACUC (protocol #2015-2294).

### 6.3 Single-cell RNA sequencing

Single cells meant for RNA processing were sorted into 96-well full-skirted Eppendorf plates that were pre-chilled at 4°C and were prefilled with 10μL of lysis buffer consisting of TCL buffer (Qiagen) supplemented with 1% beta-mercaptoethanol. Sorted plates with single-cell lysates were subsequently sealed, vortexed, spun down at 300g at 4°C for 1 minute, immediately placed on dry ice to flash-freeze the lysates, and then moved to −80°C for storage. The Smart-Seq2 protocol was performed on single sorted cells as previously described (1-3), with some modifications described below.

#### Reverse transcription

Single-cells lysates were thawed on ice for 2 minutes, then centrifuged at 3,000rpm at 4°C for 1 minute. 20μL of Agencourt RNAClean XP SPRI beads (Beckman-Coulter) was added to lysates, mixed slowly, to not introduce bubbles and subsequently incubated at room temperature for 10 minutes. The 96-well plate was then placed onto a magnet (DynaMag-96 Side Skirted Magnet, Life Technologies) for 5 minutes while covered. The supernatant was removed, and the SPRI beads were washed three times with 100μL of freshly prepared 80% ethanol, careful to avoid loss of beads during the washes. Upon completely removing ethanol after the last wash, SPRI beads were left to dry at room temperature for up to 10 minutes. Beads were resuspended in using 4μL of the following Elution Mix: 0.1μL 10μM RT primer (5’AAGCAGTGGTATCAACGCAGAGTACTTTTTTTTTTTTTTTTTTTTTTTTTTTTTTVN-3’, IDT), 1μL 10 mM dNTP (Life Technologies), 0.1μL Recombinant RNase-Inhibitor (40 U/μL, Clontech), and 2.8μL nuclease-free water. The plates were sealed and then spun down briefly, 5 seconds max to get up to 150rpm. The samples were denatured at 72°C for 3 minutes and placed immediately on ice afterwards. 7μL of the Reverse Transcription Mix was subsequently added in every well, consisting of: 2μL 5x RT buffer (Thermo Fisher Scientific), 2μL 5 M Betaine (Sigma-Aldrich), 0.09μL 1M MgCl2 (Sigma-Aldrich), 0.1μL 100μM TSO (5’- AAGCAGTGGTATCAACGCAGAGTACATrGrG+G-3’, Exiqon), 0.25 μL Recombinant RNase-Inhibitor (40 U/μL, Clontech), 0.1μL Maxima H Minus Reverse Transcriptase (200U/μL, Thermo Fisher Scientific), and 2.46μL nuclease-free water. Every well was mixed with the resuspended beads. Reverse transcription was carried out by incubating the plate at 50°C for 90 minutes, followed by heat inactivation at 85°C for 5 minutes.

#### PCR amplification and cDNA purification

14μL of PCR Mix was added in each well: 0.05μL 100μM PCR primer (5’- AAGCAGTGGTATCAACGCAGAGT-3’, IDT), 12.5μL 2x KAPA HiFi HotStart ReadyMix (KAPA Biosystems), 1.45μL nuclease-free water. The reaction was carried out with an initial incubation at 98°C for 3 minutes, followed by 22 cycles at (98°C for 15 seconds, 67°C for 20 seconds, and 72°C for 6 minutes) and a final extension at 72°C for 5 minutes. PCR products were purified by mixing them with 20μL (0.8X) of Agencourt AMPureXP SPRI beads (Beckman-Coulter), followed by a 5 minutes incubation period at room temperature. The plate was then placed onto a magnet for 6 minutes prior to removing the supernatant. SPRI beads were washed twice with 100μL of freshly prepared 70% ethanol, carefully to avoid loss of beads during the washes. Upon removing all residual ethanol traces, SPRI beads were left to dry at room temperature for up to 10 minutes. The beads were then resuspended in 20μL of TE buffer (Teknova) and incubated at room temperature for 5 minutes. The plate was placed on the magnet for 5 minutes prior to transferring the supernatant containing the amplified cDNA to a new 96-well plate. This cDNA SPRI clean-up procedure was repeated a second time to remove all residual primer dimers and resuspended in a final volume of 15μL of TE buffer. The concentration of amplified cDNA was measured using the Qubit dsDNA High Sensitivity Assay Kit (Life 7 Technologies/Thermo Fisher Scientific). The cDNA size distribution of few selected wells was assessed on a High-Sensitivity Bioanalyzer Chip (Agilent). Expected single cell cDNA quantification was around 0.5-2 ng/μL with size distribution sharply peaking around 2kb.

#### Library preparation

Library preparation was carried out using the Nextera XT DNA Sample Kit (Illumina) with indexing adapters that allow 96 single cell libraries to be simultaneously sequenced. For each library, the amplified cDNA was normalized to a 0.12-0.20ng/μL concentration range. The tagmentation reaction consisted of mixing 1.25 μL of normalized cDNA with 2.5 μL of Tagmentation DNA Buffer and 1.25 μL of Amplicon Tagment enzyme Mix. The 5 μL reaction was mixed well, spun at 3,000 rpm for 3 minutes, incubated at 55°C for 10 minutes and then immediately placed on ice upon completing this incubation step. The reaction was quenched with 1.25 μL of Neutralize Tagment Buffer and incubated at room temperature for 10 minutes. The libraries were amplified by adding 3.75 μL of Nextera PCR Master Mix, 2.5 μL of mixed indices (Nextera XT Index Kit). The PCR was carried out at an initial incubation at 72°C for 3 minutes, 95°C for 30 seconds, followed by 12 cycles of (95°C for 10 seconds, 55°C for 30 seconds, 72°C for 1 minute), and a final extension at 72°C for 5 minutes. Following PCR amplification, 2.5 μL of each library were pooled together in a 2.0 mL Eppendorf tube. The pool was mixed with 216 μL (0.9X ratio for 2.5 μl of 96 cells pooled together) of Agencourt AMPureXP SPRI beads (Beckman-Coulter) and incubated at room temperature for 5 minutes. The pool was then placed on a magnet (DynaMag-2, Life Technologies) and incubated for 5 minutes. The supernatant was removed and the SPRI beads were washed twice with 1 mL of freshly prepared 70% ethanol. Upon removing all residual ethanol traces, the SPRI beads were left to dry at room temperature for 10 minutes. The beads were resuspended in 100 μL of TE buffer and incubated at room temperature for 5 minutes. The tube was then placed back on the magnet for 3 minutes prior to transferring the supernatant to a new 1.5 mL Eppendorf tube. This SPRI clean-up procedure of the library was repeated a second time to remove all residual primer dimers, using the same approach and the final resuspension was done in 30 μL of TE buffer. The concentration of the pooled libraries was measured using the Qubit dsDNA High Sensitivity Assay Kit (Life Technologies/Thermo Fisher Scientific), and the library size distribution measured on a High-Sensitivity Bioanalyzer Chip (Agilent). Expected concentration of the pooled libraries was 30-50 ng/μL with size distribution of 300-700 bp.

### 6.4 Single-cell DNA sequencing

Single haploid cells meant for DNA processing were sorted into 96-well full-skirted Eppendorf plates either in (1) 5 μL of Cell Extraction Buffer (4.8 μL of Extraction Enzyme Dilution Buffer, 0.2 μL Cell Extraction enzyme, New England BioLabs) and processed using the PicoPlex kit, or (2) 5μL of Cell Lysis Reaction Mix (4.9 μL of Cell Lysis Buffer, 0.1 μL Cell Lysis enzyme, Yikon Genomics). Sorted plates with single-cell lysates were subsequently sealed, vortexed, spun down at 300g at 4°C for 1 minute, immediately placed on dry ice to flash-freeze the lysates, and then moved to −80°C for storage. All single-cells plates were thawed on ice for 2 minutes, then centrifuged at 3,000 rpm at 4°C for 1 minute prior to processing.

#### PicoPlex Amplification

For PicoPlex amplification, the plates were incubated at 75°C for 10 minutes, followed by 95°C for 4 minutes and held at 22°C until ready for the next steps. The pre-amplification Master Mix, consisting of 4.8 μL of Pre-Amp Buffer and 0.2 μL of Pre-Amp Enzyme Mix was added to each cell, the reaction was mixed well, spun at 3,000 rpm for 1 minute. The PCR was carried out at an initial incubation at 95°C for 2 minutes, followed by 12 cycles of (95°C for 15 seconds, 15°C for 50 seconds, 25°C for 40 seconds, 35°C for 30 seconds, 65°C for 40 seconds, 75°C for 40 seconds), and a hold at 4°C. Following the pre-amplification reaction, each well was mixed well with the Amplification Master mix, consisting of 25μL of Amplification Buffer, 34.2 μL of nuclease-free water and 0.8 μL of Amplification Enzyme Mix. The reactions were mixed well, spun at 3,000 rpm for 1 minute and incubated at 95°C for 2 minutes, followed by 16 cycles of (95°C for 15 seconds, 65°C for 1 minute, 75°C for 1 minute) and a hold at 4°C. The concentration of each cell was measured using the Qubit dsDNA High Sensitivity Assay Kit (Life Technologies/Thermo Fisher Scientific). Expected concentration of the single cell lysates was 20-50 ng/μL with size distribution of 300-1000 bp.

#### MALBAC Amplification

For MALBAC amplification, the plates were incubated at 50°C for 50 minutes, followed by 80°C for 10 minutes and held at 4°C until ready for the next steps. The pre-amplification Reaction Mix, consisting of 29 μL of Pre-Amp Buffer and 1 μL of Pre-Amp Enzyme Mix was added to each cell, the reaction was mixed well, spun at 3,000 rpm for 1 minute. The PCR was carried out at an initial incubation at 94°C for 3 minutes, followed by 8 cycles of (20°C for 40 seconds, 30°C for 40 seconds, 40°C for 30 seconds, 50°C for 30 seconds, 60°C for 30 seconds, 70°C for 4 minutes, 95°C for 20 seconds, 58°C for 10 seconds), and a hold at 4°C. Following the pre-amplification reaction, each well was mixed well with the Amplification Reaction mix, consisting of 29.2 μL of Amp Buffer and 0.8 μLamp Enzyme Mix. The reactions were mixed well, spun at 3,000 rpm for 1 minute and incubated at 94°C for 30 seconds, followed by 21 cycles of (94°C for 20 seconds, 58°C for 30 seconds, 72°C for 3 minutes) and a hold at 4°C. The concentration of each cell was measured using the Qubit dsDNA High Sensitivity Assay Kit (Life Technologies/Thermo Fisher Scientific). Expected concentration of the single cell lysates was 20-60 ng/μL with size distribution of 300-2000 bp.

### 6.5 Haplotype Phasing

#### 6.5.1 Mouse

We downloaded the combined VCF of laboratory mouse strains from The Mouse Genome project (Keane et al. 2011) and defined maternal and paternal haplotypes utilizing SNPs unique to either C57BL/6J or PWK/PHJ, respectively. For all analyses, we disregarded indels and only considered SNPs. This resulted in a total of 20,986,995 heterozygous SNPs, which overlapped 28,497 expressed genes in mouse round spermatids.

##### 10X Chromium Alignment and Haplotype Calling

We created maternal and paternal haplotypes of the two non-human primates using a combination of 10X Chromium linked read sequencing on diploid cells and sparse single cell DNA sequencing on haploid spermatid cells. We aligned the 10X Chromium reads to the *Macaca fascicularis* genome `Macaca fascicularis MacFac 5.0` from Ensembl, herein referred to as Ensembl-MF5-G, by first creating a custom reference using the longranger mkref command, and then running longranger using this reference and default parameters to generate 10X Chromium alignment data.

For Cynomolgus 1, the instrument generated 1,850,208 Gel Beads in Emulsion (GEMs) and the software mapped 819,440,960 reads for 37.2x average coverage across the genome. This resulted in 361,465 haplotype blocks with N50 length of 1.6 MB. Each block contained an average of 36 SNPs for a total of 12,758,999 heterozygous SNPs. Cynomolgus 2 10x Chromium data featured 1,889,596 GEMs that led to 812,899,614 reads mapping at an average coverage of 37.4X. It had 318,516 blocks with N50 length of 1.8 MB and an average of 40 heterozygous SNPs per block, for a total of 12,744,826 heterozygous SNPs.

##### Mature Sperm scWGS Alignment and Processing

We genotyped the haploid sperm scWGS samples using a custom pipeline. First, the paired-end reads were aligned to Ensembl-MF5 using BWA v0.7.5 (Li and Durbin 2009) using the mem option with default parameters. The resulting bam files were sorted using samtools v1.4.1 sort (Li, Handsaker, et al. 2009) and duplicates were removed using sambamba v0.6.6 (Tarasov et al. 2015). `samtools mpileup` with a bed file of the 10X Chromium identified variant positions calculated the allelic depths per heterozygous site. We then filtered the file to only include allelic depths of variant alleles. For Cynomolgus 1, this resulted in an average of 1.2M heterozygous sites per spermatid sample, for a total overlap of 3.3M sites across the 17 spermatid samples. With the 8 spermatid samples for Cynomolgus 2, we covered 3.5M total sites with an average 1.2M sites per sample at roughly 1X coverage.

##### Creating Chromosome-Length Haplotype Blocks

The final step involved stitching the the haplotype blocks generated by 10X Chromium sequencing into chromosome-length haplotypes using the haploid cell haplotypes as a guide. In the case of no recombination, the stitching is trivial and requires only a single sperm sample. Due to recombination, we used multiple sperm single cell WGS samples, and utilized a dynamic programming framework tuned to minimizing the number of recombination events to assign the chromium blocks to maternal and paternal haplotypes.

### 6.6 Allele-specific Expression Quantification

#### StringTie Transcriptome Assembly

Due to unavailability of a publicly available testes transcriptome of Cynomolgus, we created a custom cynomolgus transcriptome using single cell RNA-seq samples of round spermatid and elongating spermatid cells from the two individuals. First, we aligned the samples to Ensembl-MF5-G using STAR v2.5.3 with default parameters, and merged and sorted the bams using samtools. This resulted in 3.7 billion total reads aligned across the corpus of 480 samples. We fed the merged bam into StringTie v1.3.3 with default options except for -p 39 to indicate a large number of available threads. We compared the StringTie generated transcriptome to MacFas 5.0, a Ensembl-generated transcriptome of Macaca fascicularis using Cufflinks v.2.2.1 gffcompare, and created a dictionary to map the StringTie annotation ids back to known gene symbols.

#### RNA-Seq Processing and Alignment

To reduce allelic bias in read mapping, we used bcftools consensus to generate masked genomes, in which all bases in heterozygous positions were modified to the IUPAC character N in the reference genomes. We used STAR v2.5.3 to align the round and elongating spermatid single cell RNAseq reads, but created custom STAR genomes with either Ensembl GRCm38 or the previously described StringTie-generated transcriptomes. We utilized STAR options ‒outFilterMultimapNmax 1 to eliminate multi-mapping reads, ‒alignSJBoverhangMin 4 to force large overlap between RNA-seq reads and the genome, and ‒outSAMattributes NH HI NM nM MD XS attributes, and removed duplicated reads using sambamba. featureCounts (Liao, Smyth, and Shi 2014) was used to generate gene transcripts per million (TPM) values with options -s 0 for unstranded reads, -p for paired end reads, and -B to require both ends of the read to be mapped.

#### Generating Allele-specific Counts

To quantify allele specific expression of genes, we first assigned each heterozygous SNP to a gene using the snpEff Cingolani et al. 2012 annotate tool using custom snpEff databases. Then, after splitting the aligned RNA-seq bams into chromosome-specific BAMs, we generated the allele counts for each gene and spermatid sample combination. To avoid double-counting of reads that overlapped multiple sites, each read was only counted once in favor of either allele, and if a read matched variants on both alleles, we tagged it as a discordant read and did not utilize it for further analysis. For mouse, we average 145 allele-specific reads per gene per sample across 11,542 phaseable genes in 95 spermatid samples.

We performed an additional step to quantify allele specific counts in the non-human primate samples. Due to limited coverage across the length of an entire gene, StringTie often splits a single Ensembl gene annotation into multiple gene annotations. As such, we summed reads from separate StringTie genes overlapping known annotations. The resulting allele counts files for the monkeys are a combination of known genes annotated by Ensembl and novel genes identified by StringTie only. For Cynomolgus 1, in 187 spermatid samples, we average 122 reads per gene per sample for 8956 phased genes. For Cynomolgus 2, in 185 spermatid samples, we average 131 reads per gene per sample in 8216 genes.

### 6.7 Haplotype and Genoinformativity Inference

To study haploid-biased gene expression, we require knowledge of the underlying haplotype. We reasoned that if there was true haploid biased expression, it would be possible to infer the haplotype from the allele specific expression data. As such, we derived a model to perform both haplotype and genoinformativity inference simulataneously. Here, we first describe a model for transcript sharing across a syncytium and then extend it to a probability model for observing allele specific reads from round spermatids in a single cell RNA-seq assay.

#### Model of Genoinformative Transcripts

We begin by describing a simple model for the number of transcripts of a single gene *g* in a single cell *c*. The total number of transcripts *T* in the haploid cell is the combination of external transcripts *E* and retained transcripts *R*.

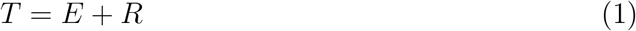

Here, external transcripts indicates transcripts that were not transcribed by the haploid cell, but rather were transported into the cell through the cytoplasmic bridge. Retained transcripts are the transcripts that were transcribed by the cell and not shared through the cytoplasmic bridge.

We can also write down the total transcripts *T* as the combination of transcripts from the maternal allele of the gene *M* or the paternal allele of the gene *P*.

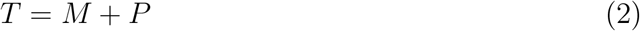

Note that we can marginalize the maternal and paternal transcripts in terms of external and retained transcripts.

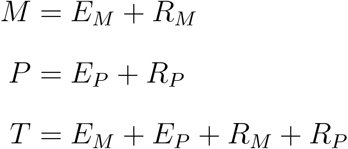

Before deriving a model for genoinformativity, we introduce two last definitions in the form of ratios. The ratio of *E*_*M*_ to *E*, or the skew of transcripts towards the maternal allele *S*, and the ratio of *R* to *T*, or the genoinformativity of the transcript.

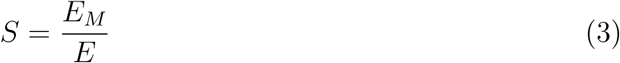

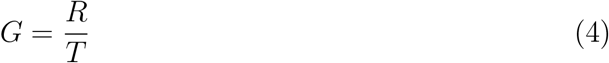

Assuming no eQTL effects, genome imprinting, technical bias, or other mechanisms for differential allelic expression, this allelic skew *S* is 0.5, i.e. the number of haploid cells that contain maternal and paternal genotype are equal and the number of transcripts transferring into the cell is equal from either allele.

#### Haploid Cell with Maternal Allele

Given the previous system and definitions, we now derive the empirical genoinformativity for a single haploid cell. Consider the case of a cell *c* having the maternal allele for the gene or haplotype *H*_*M*_. Then, we further deconvolve the total transcripts by the transcripts from the maternal allele *M* and the transcripts from the paternal allele *P*. Note that this classification is only relevant for autosomes, where it is possible to have transcripts from either chromosome in the haploid cell.

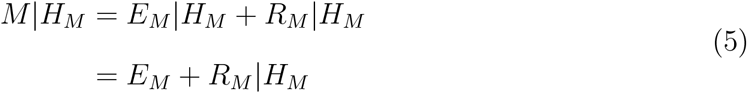

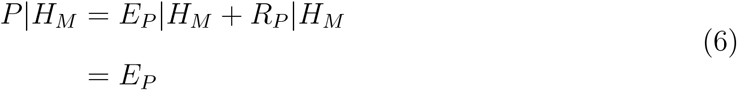

Since the cell has a maternal allele only for the gene of interest, there are no retained reads from the paternal allele. Finally, let’s express the total transcripts T in terms of the maternal and paternal transcripts.

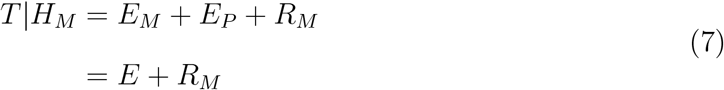

Given equation 3, 4, and 7, we can restate equation 5 as

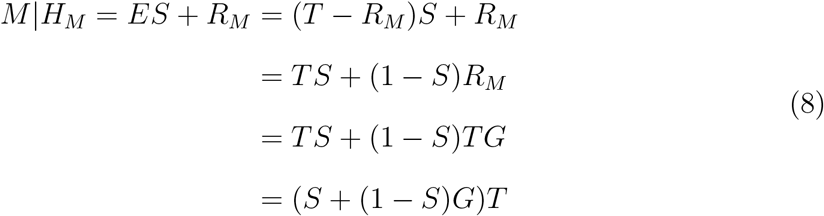

We can derive similar equations for *P*|*H*_*M*_, *M*|*H*_*P*_ and *P*|*H*_*P*_.

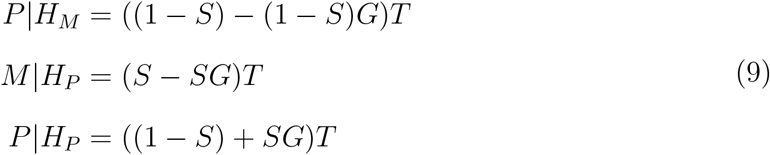

#### Probability Model for Allele-Specific Reads

We now focus our attention on developing a model for observing allele-specific reads using single cell RNA-Seq from haploid round spermatids. We derive a probability model for observing counts of alleles from the maternal allele *C*^*M*^ and paternal allele *C*^*P*^ for *N* individuals and *G* genes. Given parameters *θ*, each cell *i* and gene *j* is independent of each other and the collective probability can be written as:

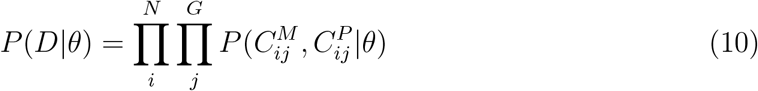

For simplicity, we will write the set of counts 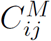 and 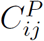 as *D*_*ij*_ where applicable. The main reason we are able to treat each set of counts independently is because we marginalize the probability over the haplotype *H*_*ij*_ of cell *i* at gene *j*.

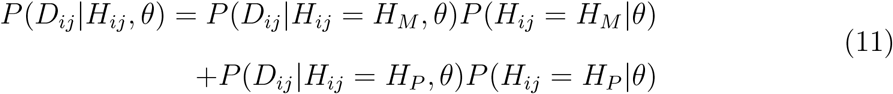

Using the above formulation, it is possible to split the inference goal into two separate sub-tasks: haplotype inference and genoinformativity inference.

#### Haplotype Inference

We use a Markov chain across a single chromosome to estimate the haplotype given a recombination rate *r*.

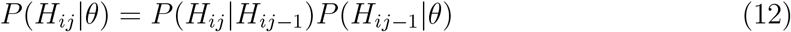

where

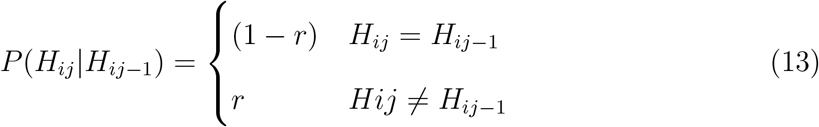

We set the initial probability of each cell’s haplotype to be equal at 0.5.

#### Genoinformativity Inference

Given the haplotype *H*_*ij*_ of cell *i* at gene *j*, the counts of the maternal and paternal allele follows from the generative model described above. Due to overdispersion in RNA-seq data, we model the counts using a beta-binomial distribution, which is specified by shape parameters *α* and *β*. In fitting the model, we only fit the shape parameter *β* and reparameterize *α* in terms of skew *S* and genoinformativity *G*. More explicitly, we can model the system as

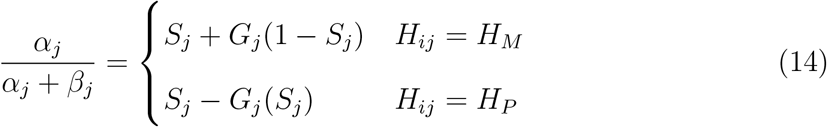

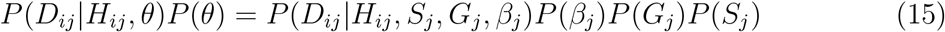

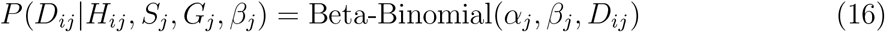

In addition to overdispersion, single cell RNA-seq data also contains high amount of allelic dropout and amplification of a single molecule. To alleviate the impact of allelic dropout on estimates of genoinformativity, we introduce a Zero-and-N-inflated Beta Binomial distribution parameterized by an additional variable *ζ*_*j*_ which defines the probability of allelic dropout for the gene.

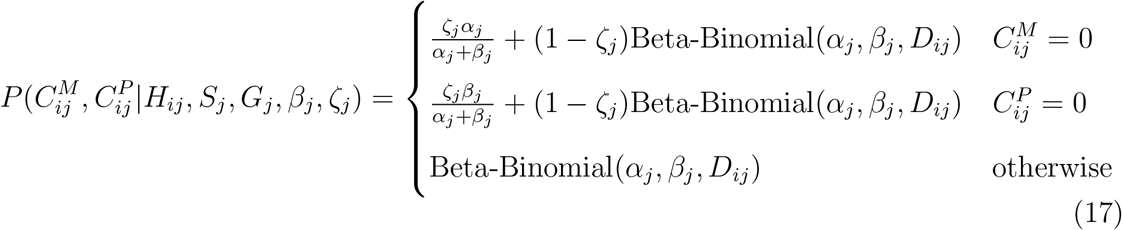

#### 6.7.1 Implementation

##### Haplotype Inference

Unfortunately due to inherent noise in the system and the cost of sampling the aforementioned Markov chain, we do not compute the Markov chain for each gene independently. Instead, we bin the genes into buckets *B* and perform a similar inference task with each bucket *k*. Each bucket on average contained 10 genes in our fits.

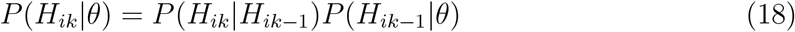

We also used a fixed recombination rate *r* for each cell and each chromosome with the assumption that a cell would have on average 0.5 recombination events per chromosome.

##### Genoinformativity Inference

Instead of learning the parameter *S*_*j*_ for each gene, we use a empirical estimate of *S*_*j*_ derived from dividing the number of *H*_*M*_ reads for a gene *j* by the total number of reads for that gene across all cells. We also tested using the mean of the empirical *S*_*ij*_ derived from each cell separately, and did not notice large differences in the model fits.

##### Priors

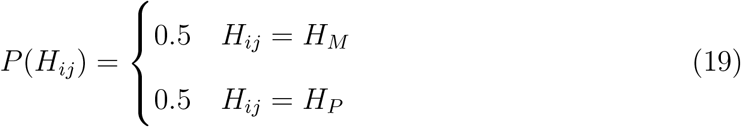

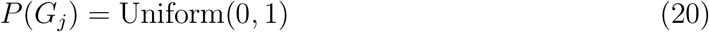

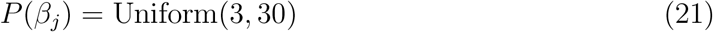

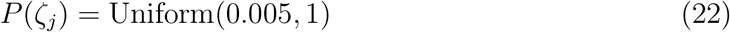

##### Two-stage Fitting

For computational efficiency, we split the inference task into two stages. In the first stage, we fit both the haplotype and genoinformativity inference steps for highly expressed genes (TPM > 20). Then, in the second stage, we only performed genoinformativity inference using fixed haplotype probabilities. We used the mean posterior of the haplotypes from the first stage, and interpolated the probability for genes that were unique to the second stage. There was 99% correlation between the posterior mean of the genoinformativity values, indicating low variance in the posterior haplotype distributions and high confidence in haplotype inference.

##### Samplers

We used PyMC3 (Salvatier, Wiecki, and Fonnesbeck 2016) as the framework for sampling the model. For the haplotype sampling, we used a Categorical Gibbs Metropolis sampler. All the other parameters were sampled using the No U-turn Sampler (NUTS) with a target accept probability of 0.8. We sampled the model for 5000 steps with two separate chains and used the last 500 steps for estimating the posterior distribution across the 2 chains.

#### 6.7.2 Sex chromosome GIMs

Mouse gene-level transcripts per million (TPM) values were collected for all genes in all spermatids using all RNAseq reads, not only allele-informative reads. For each gene, a loess regression was used to fit its log2 expression across the diffusion pseudotime with a pseudocount of 1 TPM, using the R loess function with a gaussian function family and 0.75 span. The residuals from this fit were then used to calculate pairwise Spearman correlations between all sex chromosome genes. Pairwise correlations were hierarchically clustered using the complete linkage method, with the results visualized in heatmaps. A cutoff height of 6 was empirically found to split the data into three clusters: a distinct X cluster, an anti-correlated distinct Y cluster, and a mixed X and Y cluster with no strong correlation patterns. Genes in the first two clusters were considered potential GIMs. We calculated the mean pairwise Spearman correlation between pairs of potential GIMs, with the sign reversed for genes in opposite clusters. Genes with a mean pairwise correlation of greater than 0.05 (roughly the median value over potential GIMs) were selected as putative sex chromosome GIMs.

### 6.8 GIM classification

To classify each gene as a “Confident GIM”, “Confident Non-GIM”, or “Remaining Gene”, we fit the Bayesian model to shuffled data, and compared the posterior distributions for *H*_*i*_*k*, *G*_*j*_, and *β*_*j*_ between real and shuffled data. We utilized two main shuffling methods: complete shuffle and cell-label shuffle for each chromosome independently. The complete shuffle shuffled the allele counts randomly across the population of cells and genes. For the cell-label shuffle, the allele counts were randomized across the cells, but the distribution of counts in a gene remained the same. We trained our Bayesian HMM using the same default parameters and priors as the real data, and compared the model fits. Since *β*_*j*_ can capture both the variance of single cell rna-seq as well as the variance in genoinformativity, we created a new measure γ_*j*_ as an alternative measure of genoinformativity that combines both posterior mean estimates of *G*_*j*_ and *β*_*j*_.

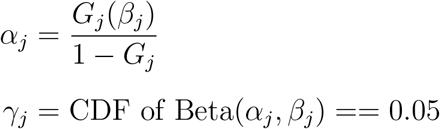

To reflect the confidence of the haplotype fits *H*_*ik*_ across all *n* samples, we also created an aggregated measure, fraction of poor haplotypes 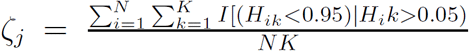, which reflected the proportion of haplotypes that a posterior mean haplotype probability less than 0.95 for either the maternal or paternal haplotypes.

We performed a grid search across thresholds for highest posterior density (hpd) evaluated at 5% and 95% for genoinformativity *G*_*j*_ and γ_*j*_and*ζ*_*j*_, which controlled the eFDR at 10% for confident and non-confident gims compared to the shuffled control. For a particular gene, the thresholds for a “Confident GIM” are: hpd 5% of genoinformativity > 0.025, hpd 95% of genoinformativity > 0.2, γ_*j*_ > 0.025, fraction of poor haplotypes < 0.4 For “Confident Non-GIMs” are restricted to hpd 95% of genoinformativity < 0.2. Genes that fall outside these bounds were considered “Remaining Genes”.

### 6.9 GIM characterization

#### Expression-matched control selection

The expression trajectory across spermiogenesis was first tabulated for each gene by cross-referencing the log2 of the TPM expression level (with a pseudocount of 1 and complete dropout considered a zero) against the diffusion map pseudotime value for each cell (i.e. the first dimension of the map). To reduce noise at the single cell level, a smoothed loess fit was used as the expression trajectory (fit using default parameters for the `R loess` function).

Next, all confident non-GIMs expressed in spermatids were considered as controls for all GIMs. Pools of controls were first reduced for each GIM based on two hard filters: first, all genes were equally distributed into 5 bins based on their dropout rates; second, the slope of a linear fit to the expression trajectory was required to differ by no more than 0.2. This helped to control for any confounders resulting in oversampling, as well as large expression changes in a small number of cells, generally in the extreme early or late part of the trajectory.

For each GIM, all non-GIMs remaining in its pool were ranked by their mean squared difference in log2 expression level, and the top 20 were selected as mock GIM controls, whose ranks were then scrambled. This resulted in 20 control sets of mock GIMs having similar dropout rates, slope of expression trajectory, and low difference in expression trajectory. For analyses limited to protein coding genes, control selection was performed again with both the GIMs and the control pools limited to protein coding genes.

For the cynomolgus samples, the expression trajectories were averaged across the two individuals. Where stringtie genes overlapped with Ensembl annotations, the aggregated expression for the Ensembl annotation was used for both GIMs and controls. A gene was considered a GIM if it was called as a confident GIM in either individual, and was considered as a non-GIM if it was called as a confident non-GIM in either indivdual. The rare genes having conflicting calls in each individual were excluded from these analyses. Human GIMs and non-GIMs were inferred from homologous cynomolgus annotations with homology defined as having the same Ensembl gene symbol (i.e. standard gene name).

For spermatid-expressed non-GIM controls, control sets were selected from all confident non-GIMs randomly, without filtering for dropout bin or expression trajectory fit.

#### Gene Ontology

Mouse gene ontology annotations were downloaded from Ensembl Biomart with the Ensembl Genes 93 / GRCm38.p6 annotation dataset. The mean and standard deviation of number of GIMs expected with each annotation was calculated based on the 20 control sets. Nominal probabilities were then calculated using the normal distribution, and multiple testing was corrected using the Benjamini-Hochberg method to result in false discovery rates. GO terms were considered significant if they had at least 20 GIMs, an FDR ≤ 0.001 and a moderated log2 enrichment (using a pseudocount of 5) of at least 0.5.

For COMPARTMENTS comparisons, fewer controls had at least one annotation than GIMs, which could artificially inflate significance for individual categories. Therefore, we performed an additional normalization for the expected number of GIMs with an annotation. The number of controls in a set having a GO annotation was converted to a fraction out of those have any annotation, and then multiplied by the number of GIMs to yield the total number expected with each annotation specifically. Otherwise the enrichment analysis was the same as for the GO analysis above.

#### 3’ UTR motifs

Only protein-coding genes were considered. The 3’ UTR annotations of GIMs and their controls were taken from the highest expressed Ensembl transcript in spermatids. UTRs annotated as less than 7 nucleotides in length were discarded. All 20 sets of control UTRs were combined into a single background set, allowing duplicates. AME, a tool from the MEME suite, was run with default parameters using a motif database comprised of the CISBP-RNA and Ray2013 mouse and human sets provided by MEME. The enrichment search was performed using GIMs as foreground and the combined control set as background, with foreground and background switched for the depletion analysis.

For candidate RNA-binding proteins, only those with a maximum TPM of 10 at any point in the loess-smoothed expression trajectory were considered. Enrichments were considered significant at an E-value cutoff of 0.01. Motifs having the same IUPAC consensus were merged into a single result.

#### Selective sweeps

Candidates for mouse selective sweeps were taken from Staubach et al. 2012. Sweep regions in any population were considered. All candidate genes within 600kb of each other were collapsed into a single region. For GIMs or each control set, the number of regions having at least one overlapping gene was counted. For a p-value of this difference, the mean and standard deviation of the control sets was used to generate a one-sample t-test.

Candidates for human selective sweeps were taken from Refoyo-Martınez et al. 2019; Schrider and Kern 2016; Ferrer-Admetlla et al. 2014; Cheng, Racimo, and Nielsen 2019; Munch et al. 2016, with selective sweep regions as in each paper. In cases in which the paper predicted selective sweep regions but did not annotate associated genes, all genes overlapping the regions were considered selective sweep candidates. Otherwise this analysis was as in the mouse.

Direct testing for human selective sweeps was performed using statistics from Pybus et al. 2014 based on analysis of the 1000 genomes project data. To help control for differences in gene length, the median score overlapping the 3’ UTR was used to represent the gene. The “best” score for each gene was taken across each population, where “best” signifies the raw score most in favor of a selective sweep for that score. Selective sweep candidates were defined as any where the score was at least 3 standard deviations beyond the mean in this direction. The number of GIM sweep candidates was compared to background expectation of the mean and standard deviation among the 20 control sets.

#### Testis-specific paralogs

Paralog and tissue-specificity data were taken from Guschanski, Warnefors, and Kaessmann 2017. Testis-specific paralogs were defined as those with a “Tissue specificity” (as defined by the paper) of at least 0.90.

#### Alternative splicing

Mouse alternative splicing was taken from events in VastDB with quality greater than zero and testis specific was defined as a difference in PSI of at least 50 between testis and the median PSI across all other tissues.

Human isoform expression was taken from the GTEX consortium (file `GTEx Analysis 2016-01-15_v7_`) Since individual isoform estimates can be unstable, we considred subsets of isoforms that are expressed higher in testis. Each transcript was ranked by the difference between testis isoform usage (i.e. ratio of transcript TPM to gene TPM in that tissue) to the median tissue isoform usage across other tissues. The maximum of the cumulative sum of excess isoform usage in testis was counted as the testis specificity (testis isoform usage minus other tissue isoform usage). A cutoff of 0.5 was considered testis-specific (equivalent to 50 PSI).

#### Late translation

Translation data was taken from Iguchi, Tobias, and Hecht 2006 (GSE4711 on GEO). Translation efficiencies were calculated as the median across replicates of the fold change from polysome to RNP samples. Genes were defined as having specific late translation if they were in the bottom quartile of this score at day 22 (which is depleted for late spermiogenesis), and the top quartile with respect to fold-change increase in translation efficiency between day 22 and adult mice. For each of the functional readouts of GIMs (e.g. selective sweeps), we compared the fraction of GIMs in that category that were specifically late translated to those that were not in that category (i.e. not functional candidates by that measure).

## Supporting information

Zip of all supplementary tables

---

**Supplemental figure 1.**
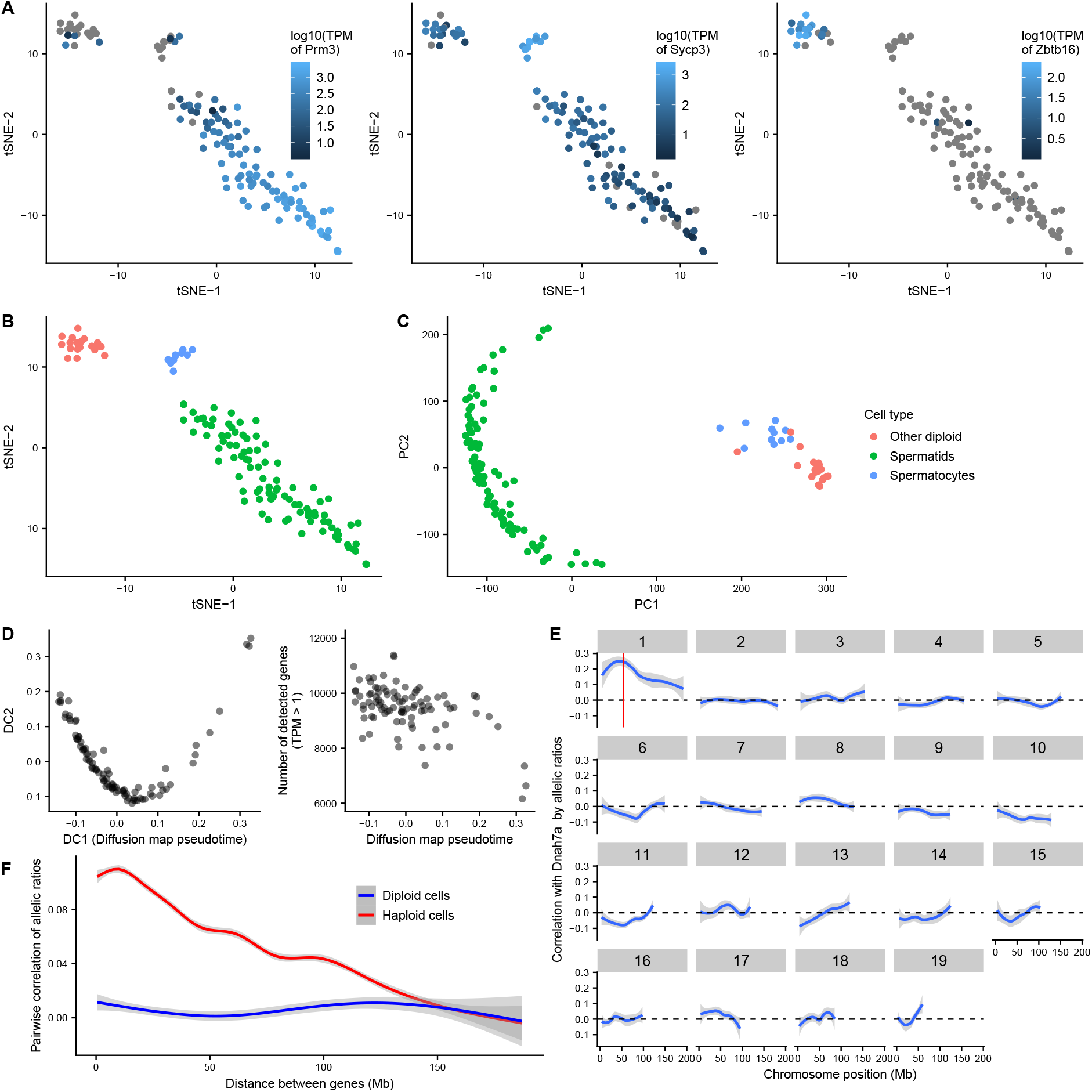
Single cell RNAseq of haploid spermatids identifies chromosome-scale correlations in allelic bias. **A)** t-Distributed Stochastic Neighbor Embedding (tSNE) dimensionality reduction for single testis cells enriched for haploid cells. Expression levels in transcripts per million (TPM) are visualized for markers of haploid spermatids (*Prm3*), spermatocytes (*Sycp3*), and spermatogonia (*Zbtb16*). **B)** Cell type annotations based on the above marker genes. **C)** Principal component analysis confirming the tSNE result, showing that all haploid spermatids were strongly distinct from diploid cells. **D)** Left: first two dimensions of diffusion map of haploid spermatids showing the first dimension captured the developmental stage well. Right: Number of genes detected per cell against the first diffusion map dimension (diffusion map pseudotime), showing a decline in those at the latest developmental stage. **E)** Illustration of chromosome-length allelic expression correlation. For one gene on chromosome 1, *Dnah7a* (located at the red line), pairwise correlation of allelic expression ratio was calculated for every gene. Plotted is a loess-smoothed average across each chromosome. Only on chromosome 1 near the *Dnah7a* locus is there a substantial average correlation. **F)** Summary of chromosome-length allelic expression correlations. For each gene, pairwise correlations of allelic expression ratios with all genes on the same chromosome were calculated. The mean correlation in haploid cells or diploid cells across all genes is plotted as a loess-smoothed average. A substantial mean correlation exists for nearby genes in haploid but not diploid cells, and decreases gradually across tens of megabases.

**Supplemental figure 2.**
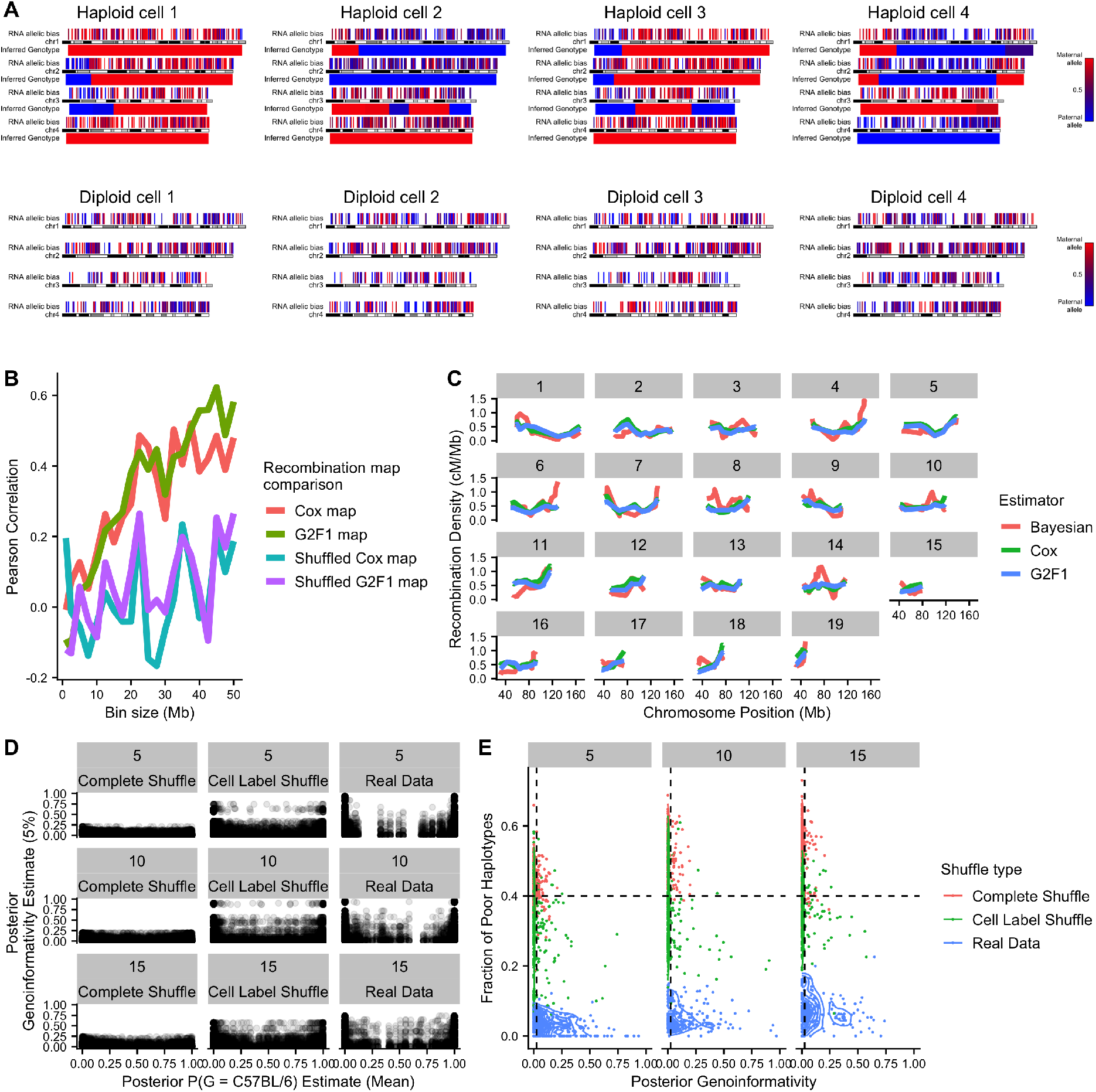
Joint inference of genotype and genoinformativity. **A)** Visualization of allelic bias in the first four chromosomes of randomly selected haploid cells and randomly selected diploid cells. Each expressed gene is represented as a vertical line with color representing its allelic ratio (red for more maternal allele, blue for more paternal). Below each chromosome is the genotype automatically inferred by our Bayesian method. **B)** Correlations between inferred recombination densities and two published mouse recombination maps (Cox et al. 2009; Liu et al. 2014) or corresponding controls with recombination densities shuffled between all bins. **C)** Recombination densities across each chromosome (calculated over a 20Mb window) implied by the Bayesian recombination frequencies or for each of the two published recombination maps. **D)** Inferred genotype and genoinformativity for real haploid data and two shuffle types: one permuting both gene and cell labels (complete shuffle) and one permuting only cell labels. Each point is a gene/cell pair, with genotype estimate (x-axis) being a property of the specific gene in a specific cell, and 5% lower bound of genoinformativity (y-axis) being a property of the gene (constant across cells). Three representative chromosomes are plotted (5, 10, and 15). Real data more often have confident genotype estimates and high genoinformativity (upper left and upper right of graph). The cell label shuffle is quite conservative because the genotype structure is maintained, and only the genoinformative expression is randomized. **E)** Summary of the data from (D) illustrating thresholds for calling confident GIMs (dashed lines). Each point is a gene, with poor haplotypes defined as those with less than 95% probability of a genotype. 5% lower bound of posterior genoinformativity probability is plotted on x-axis.

**Supplemental figure 3.**
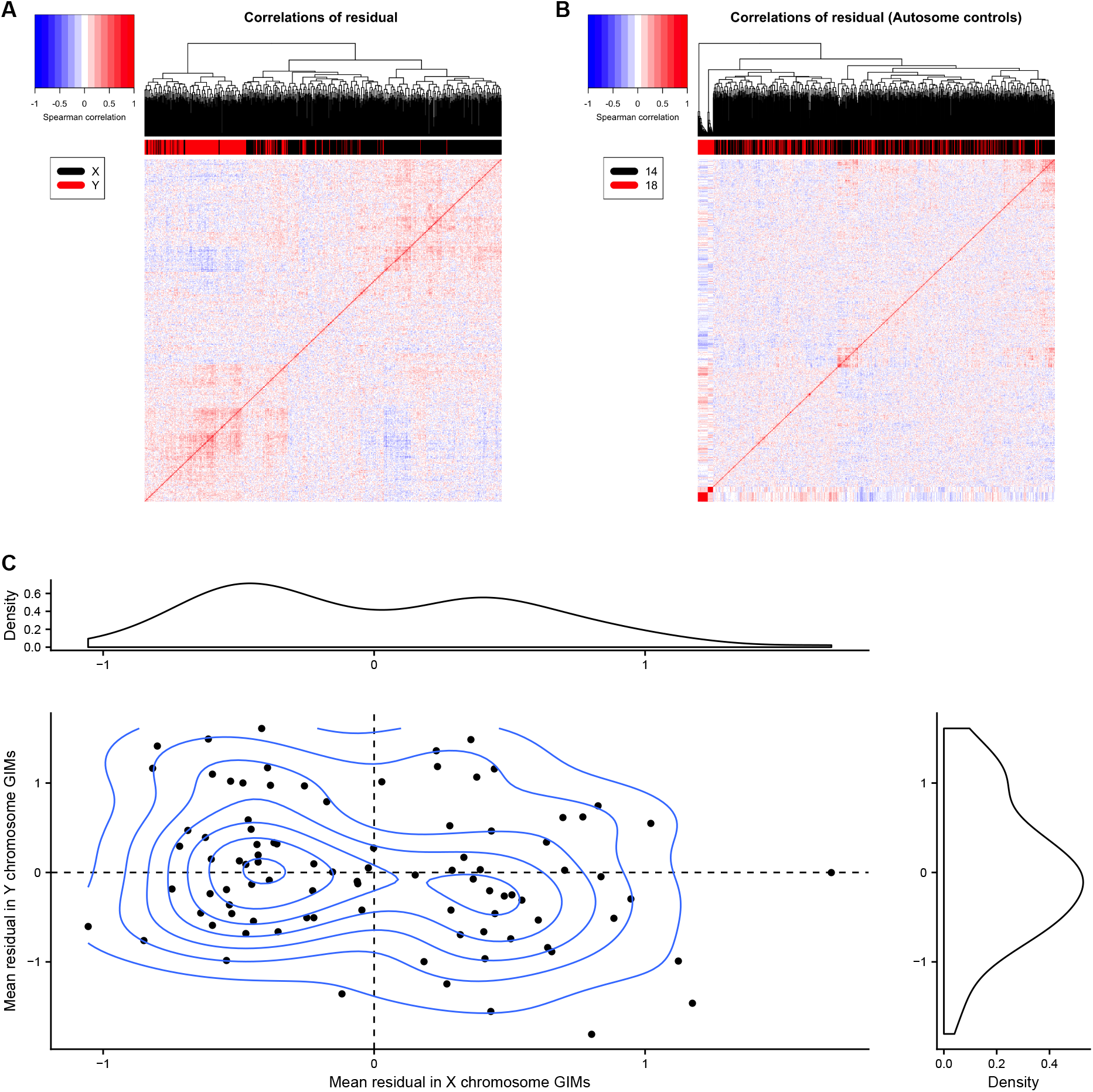
Sex chromosome GIMs. **A)** Heatmap of pairwise correlations of sex chromosome genes. Correcting for developmental stage (fitting the expression to the diffusion pseudotime position), the residuals of the log expression levels are correlated between all pairs of sex chromosome genes. Two anticorrelated clusters appear, one principally on the X chromosome (black lines above the heatmap), one principally on the Y chromosome (red lines above the heatmap). **B)** Heatmap of pairwise correlations as in (A), but for autosomal control chromosomes with similar numbers of spermatid-expressed genes (chromosomes 14 and 18). No similar broad clusters appear. **C)** Cells have bimodal expression of putative X chromosome GIMs. For each cell, the mean residual log expression across putative X GIMs and Y GIMs is plotted, with density contours. Density plots on the margins show the kernel density of the mean residual for X GIMs (top) and for Y GIMs (right). Most cells have either a high or a low average expression of X chromosome GIMs, but not intermediate. Cells that have high X GIM expression tend to have lower expression of Y GIMs, and vice versa.

**Supplemental figure 4.**
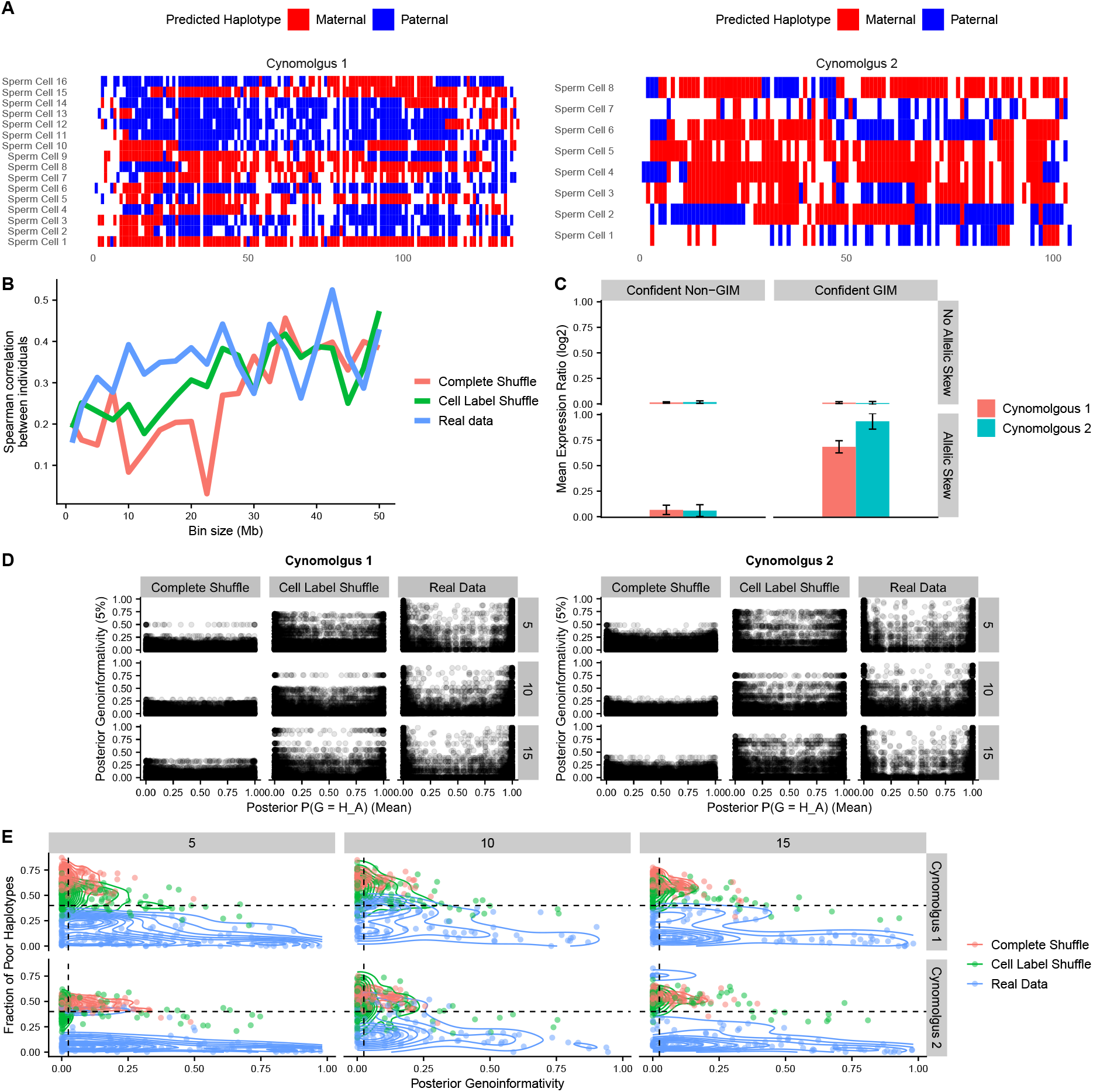
Cynomolgus primate genotype and genoinformativity inference. **A)** Single cell DNA sequencing data is displayed as phasing blocks called by the 10x Chromium pipeline for chromosome 1. Blocks are assigned to parental chromosomes based on the single cell sequencing data using the algorithm described in the methods section. The resulting patterns show 1-2 recombinations per cell with very few discordant (incorrectly assigned) blocks. **B)** Spearman correlation between recombination densities inferred for the two individuals. Shuffled data showed lower correlations at low to moderate bin sizes. **C)** Summary of expression differences (log2 ratio of genotype concordant with skew to discordant) in all genes in each of the four combinations listed. Only with both allelic skew and GIMs is there an expression difference between cells of differing genotypes, matching the results in mouse. **D)** Inferred genotype and genoinformativity for real haploid data and two shuffle types: one permuting both gene and cell labels (complete shuffle) and one permuting only cell labels. Each point is a gene/cell pair, with genotype estimate (x-axis) being a property of the specific gene in a specific cell, and 5% lower bound of genoinformativity (y-axis) being a property of the gene (constant across cells). Three representative chromosomes are plotted (5, 10, and 15). Real data more often have confident genotype estimates and high genoinformativity (upper left and upper right of graph). The cell label shuffle is quite conservative because the genotype structure is maintained, and only the genoinformative expression is randomized. **E)** Summary of the data from (D) illustrating thresholds for calling confident GIMs (dashed lines). Each point is a gene, with poor haplotypes defined as those with less than 95% probability of a genotype. 5% lower bound of posterior genoinformativity probability is plotted on x-axis.

**Supplemental figure 5.**
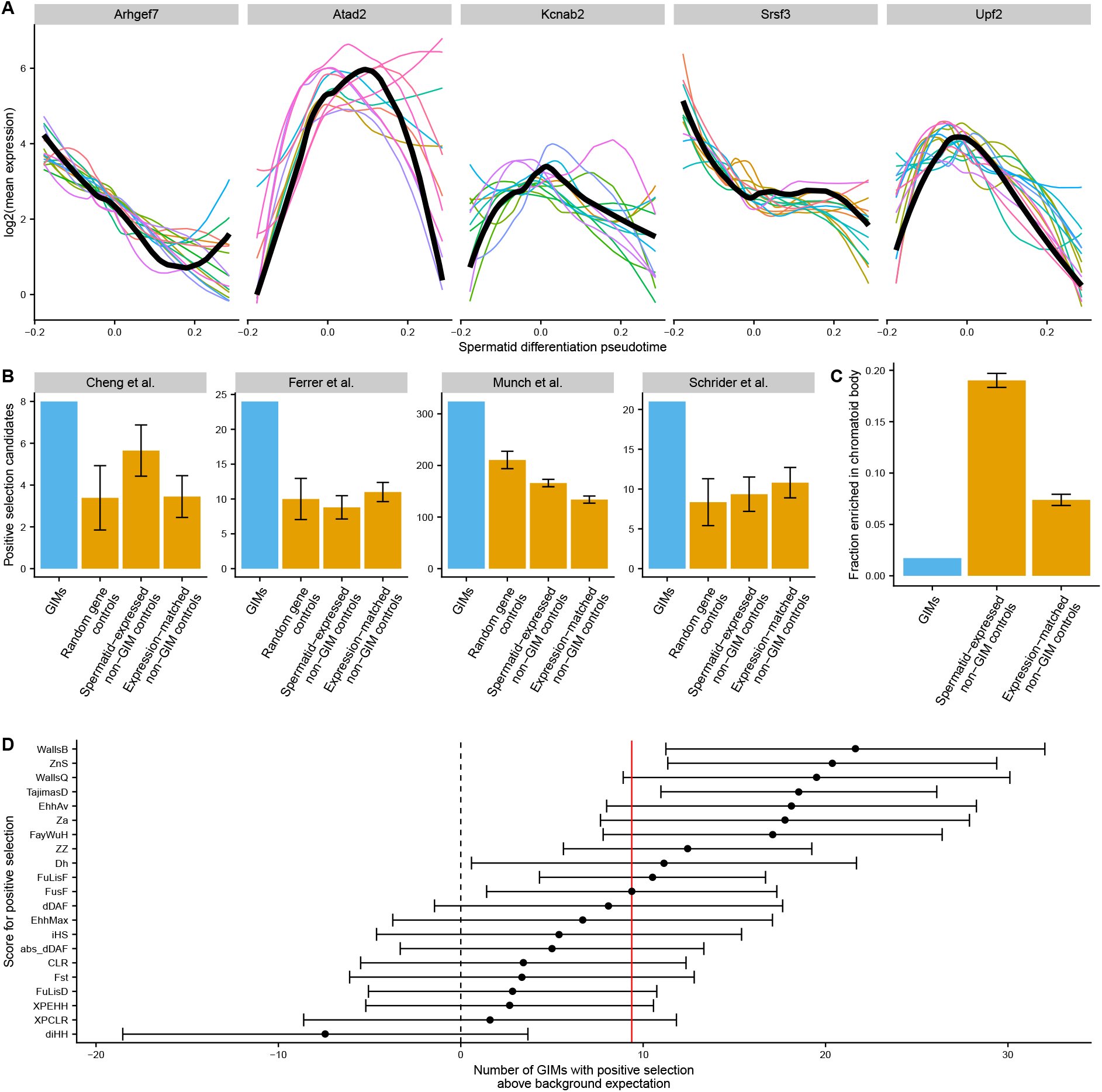
GIM functional characterization. **A)** Illustration of expression-matched control selection for representative GIMs. Thick black lines represent log2 of the loess fit of the expression (in TPM) of GIMs across the spermatid differentiation diffusion pseudotime. Colored lines represent the same loess fit for the 20 genes selected as controls for this gene based on their expression pattern and dropout rate. **B)** The number of positive selection (selective sweep) candidates from several publications (Schrider and Kern 2016; Ferrer-Admetlla et al. 2014; Cheng, Racimo, and Nielsen 2019; Munch et al. 2016) overlapping GIMs or several types of controls. Error bars represent the mean standard deviation over the 20 control sets of mock GIMs. GIMs are enriched for selective sweeps in all cases (p < 0.0276, p < 1.01 × 10^−6^, p < 9.65 × 10^−6^, p < 8.60 × 10^−6^, respectively). **C)** The fraction of genes overlapping the genes annotated as enriched in the chromatoid body (Meikar et al. 2014) overlapping with each gene category. Bars represent mean standard deviation over the 20 control sets of mock GIMs. **D)** Enrichment for GIMs in positive selection candidates based on raw scores for positive selection calculated based on 1000 genomes project data. The background expectation was calculated using the expression-matched non-GIM control set, and error bars represent the mean *±* twice the standard deviation of these controls.

